# Wiz regulates clustered protocadherin genes by restricting CTCF/cohesin loop extrusion in a genomic-distance biased manner

**DOI:** 10.64898/2026.03.02.708952

**Authors:** Tianjie Li, Jingwei Li, Leyang Wang, Haiyan Huang, Qiang Wu

## Abstract

Zinc finger (ZF) proteins (ZFPs) constitute the largest family of transcription factors in mammals; however, their regulatory mechanism remains largely elusive. Here we propose COP (C2H2-ZFP occupancy predictor), a deep learning framework that integrates DNA sequence information with protein primary and secondary features to systematically predict ZFP genomic enrichments. Applying COP to the mouse clustered protocadherin (*cPcdh*) gene locus, we identified dozens of C2H2-ZFPs potentially involved in CTCF-mediated gene regulation with Wiz having the highest number of 12 ZFs. We confirmed Wiz enrichments at all of the CBS (CTCF-binding site) elements across the *Pcdh* clusters by Myc-tagging the endogenous *Wiz* gene. Genetic experiments revealed significant increases of expression levels of the *cPcdh* genes upon *Wiz* deletion in both neuronal cells *in vitro* and in mouse brain *in vivo*. Finally, integrated ChIP-seq, RNA-seq, and 4C-seq analyses demonstrated that Wiz regulates CTCF/cohesin occupancy and long-range enhancer-promoter contacts in a genomic-distance biased manner. Together, these findings have interesting implications regarding molecular mechanism of C2H2-ZFPs in *cPcdh* gene regulation.

## Introduction

The *cPcdh* locus encodes dozens of cadherin-like cell adhesion proteins via stochastic and monoallelic promoter choice combined with alternative splicing [1-7]. The mouse *cPcdh* locus spans nearly 1 Mb and comprises three consecutive clusters of *α, β*, and *γ*. Both the *Pcdh α* and *γ* clusters are organized into a 5’ variable region with a tandem array of large exons and a 3’ constant region that contains a single set of common exons (Fig 1A). Each 5’ variable exon encodes a signal peptide, followed by six cadherin ectodomains, the transmembrane domain, and a membrane-proximal cytoplasmic region. The 3’ shared constant exons encode a membrane-distal C-terminal cytoplasmic region. Each 5’ variable exons is spliced to the respective set of downstream 3’ constant exons, generating 14 *Pcdhα* isoforms (12 alternate and 2 c-types) and 22 *Pcdhγ* isoforms (12 a-type, 7 b-type, and 3 c-type). The *Pcdhβ* cluster contains only 22 variable exons each encoding a *Pcdhβ* isoform [1, 8]. Each variable exon, except for *Pcdh αc2, β1, γc4*, and *γc5*, contains a promoter CTCF-binding site (CBS) element (Fig 1A). These promoter CBS (pCBS) elements are paired with CBS within a downstream distal super-enhancer to form CTCF/cohesin-mediated enhancer-promoter (E-P) chromatin loops which determine stochastic isoform choice [6, 8-11]. Each alternate variable exon of the *Pcdhα* cluster also contains an exonic CBS (eCBS) element that reinforces the chromatin loop via a “double clamping” mechanism using two pairs of CBS elements with opposite orientations [10]. This pairing is achieved via CTCF anchoring or stabilizing cohesin “loop extrusion” from downstream distal super-enhancers to upstream variable promoters in the opposite orientation of tandem arrays of forward CBS elements [7, 12, 13].

**Fig 1.**
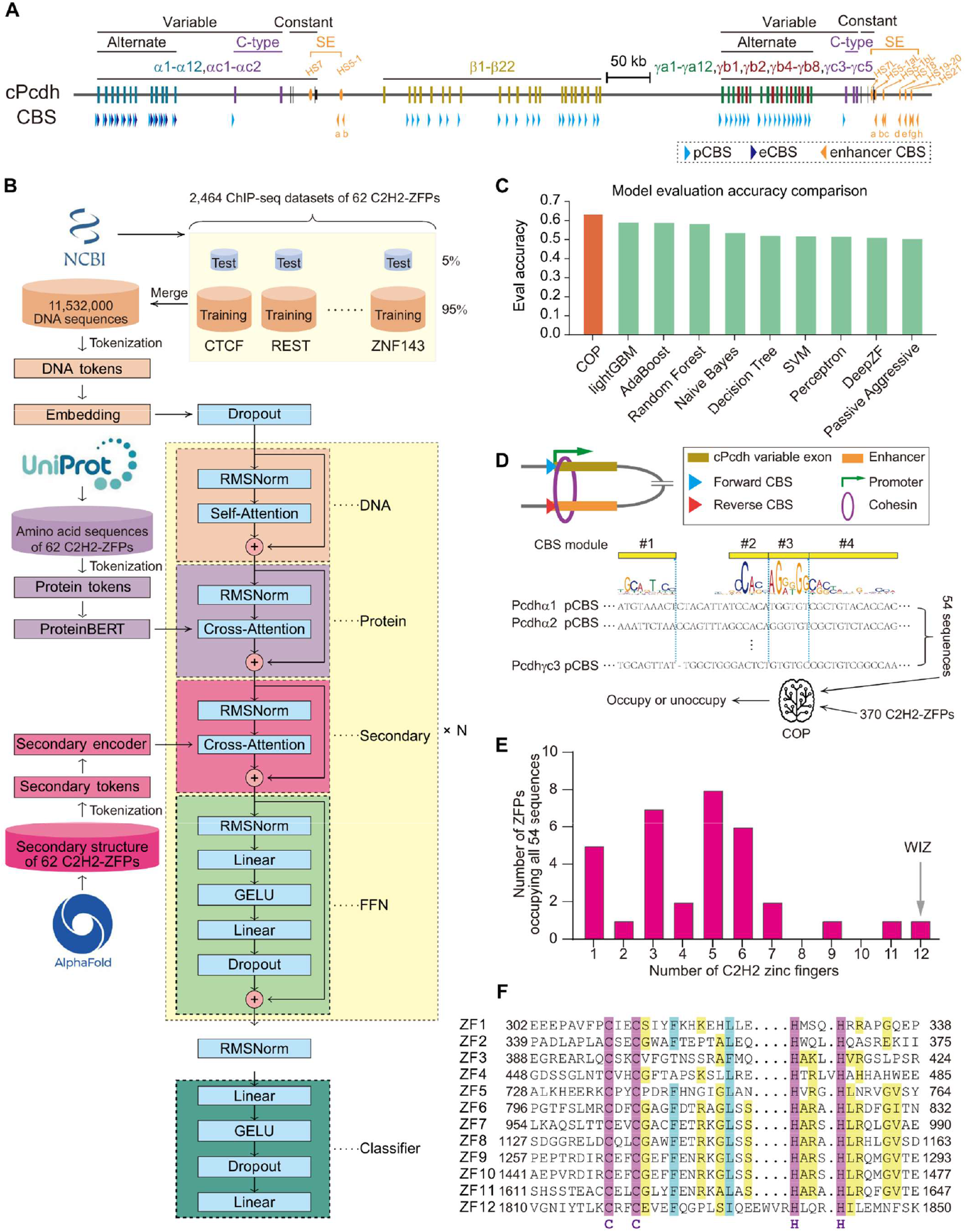
Prediction of C2H2-ZFPs localized at *cPcdh* promoter CTCF sites by COP. (**A**) Genomic organization of the mouse *cPcdh* locus. The mouse *cPcdh* locus comprises three consecutive clusters of *α, β*, and *γ*. The *α* and *γ* clusters each contain a variable region of tandem-arrayed alternate and C-type variable exons, and a constant region consisting of three constant exons. Each variable exon can be cis-spliced to the downstream set of constant exons, giving rise to 14 *Pcdhα* and 22 *Pcdhγ* isoforms. In contrast, the *β* cluster contains only variable exons and lacks a constant region, encoding 22 *Pcdhβ* isoforms. Each variable exon harbors a promoter CBS element (pCBS), with the exceptions of *Pcdh αc2, β1, γc4*, and *γc5*. Moreover, each variable exon of the *Pcdhα* cluster harbors an exonic CBS element (eCBS). The *Pcdh α* super-enhancer (*HS7* and *HS5-1)* and *Pcdh βγ* super-enhancer (*HS7L, HS5-1aL, HS5-1bL*, and *HS18-21*) are highlighted in orange. CBS, CTCF binding site; eCBS, exonic CBS; HS, hyper sensitive site; kb, kilobases; pCBS, promoter CBS; Pcdh, protocadherin; SE, super-enhancer. (**B**) Schematic of the COP model. DNA embeddings are first processed by a self-attention layer to capture intra-sequence dependencies. They then undergo two sequential cross-attention steps, first with protein primary sequence embeddings, followed by secondary structure embeddings, to integrate protein features. The fused representations are refined by a feedforward network (FFN). This entire process is repeated N times to enable hierarchical feature learning. Finally, the resulting output is fed into a classifier. COP, C2H2-ZFP occupancy predictor; FFN, feedforward network; GELU, Gaussian error linear unit; RMSNorm, root mean square layer normalization. (**C**) Benchmark comparison with existing models. The COP prediction accuracy (orange) outperforms that of other approaches. (**D**) Regulatory architecture of the *cPcdh* locus. CTCF facilitates gene activation by bringing promoters into proximity with distal enhancers by anchoring cohesin-mediated chromatin loop extrusion at forward-oriented promoter and reverse-oriented enhancer CBS elements. To predict ZFPs that cooperate with CTCF in this regulatory mechanism, the repertoire of 54 pCBS elements of *cPcdh* and 370 C2H2-ZFPs were analyzed with COP. (**E**) COP predicts Wiz as a top candidate with the highest number of 12 ZFs. (**F**) Amino acid sequence alignment of the twelve C2H2 ZFs of the mouse Wiz. The conserved cysteine (C) and histidine (H) residues that coordinate the zinc ion are highlighted in magenta.

The eleven zinc finger (ZF) domains of CCCTC-binding factor (CTCF) play a key role in antiparallel recognition of diverse *cPcdh* CBS elements for cohesin directional “loop extrusion” [6, 7, 12, 14]. Recently, several other ZF proteins (ZFP) have been implicated in regulating the *cPcdh* gene expression via directional recognition of diverse DNA elements, such as the repressor element-1 silencing transcription factor (REST) or neuron-restrictive silencer factor (NRSF) [15], ZNF143 [16], and ZNF274 [17]. They utilize different combinations of ZF domains, each typically interacting with a DNA triplet to bind to a variety of regulatory elements in a sequence-specific manner. In mammals, there are approximately ∼800 C2H2-ZFPs that constitute the largest family of DNA-binding transcription factors [18, 19]. Most of them contain clustered ZF domains, with a great variety of residues both within and between ZFs, which posing a major challenge to accurately define their DNA-binding specificity. Based on ZFP ChIP-seq (Chromatin immunoprecipitation followed by high-throughput sequencing) and ChIP-exo (ChIP with lambda exonuclease digestion) *in vivo* as well as *in vitro* binding datasets, several computational models have been developed for prediction of their binding sites [19-23]. For example, DeepZF mainly utilizes zinc finger polypeptide and target sequences as inputs to predict its DNA binding [24]. However, the biological function of ZFP enrichments at genomic sites remains challenging owing to both their flexible direct recognition of DNA motifs and indirect interacting with co-occupied proteins or even RNAs.

The widely interspaced zinc finger-containing protein (Wiz), which contains 12 dispersed ZF domains (ZF1-12), functions as a regulator of CTCF/cohesin-mediated chromatin loops [25]. Wiz was originally identified in the mouse brain as two isoforms: a long ZF1-11 and a short ZF6-11 [26]. The association of Wiz with the G9a/GLP histone methyltransferase complex suggests a transcriptional repression function [27, 28]. Recently, its repressive role in fetal hemoglobin (HbF) gene expression was uncovered through a chemical screen, revealing *Wiz* as a promising therapeutic target for sickle cell diseases (SCD) [29]. Finally, Wiz has been implicated in transcriptional insulation and is essential for maintaining embryonic stem cell identity [30]. Here we developed a C2H2-ZFP occupancy predictor (COP) using attention mechanisms to jointly model the relevance of ZFP amino acid sequences, secondary structures, and target DNA sequences. COP inferred that Wiz has the potential to bind all of the pCBS elements within the *Pcdh* gene clusters. By deleting *Wiz* in the neuroblastoma Neuro-2a (N2a) cells in vitro and in mouse brain in vivo, combined with integrated RNA-seq, ChIP-seq, and 4C-seq analyses, we found that Wiz impairs CTCF binding, restricts chromatin cohesin, and suppresses *cPcdh* gene expression in a genomic-distance biased manner.

## Results

### Prediction of C2H2-ZFPs localized at promoter CBS elements of *cPcdh* by COP

To systematically predict C2H2-ZFP localization at specific DNA sequences, we developed a deep neural network model named COP, which comprises separate encoding modules for DNA sequences, protein sequences, and protein secondary structures (Fig 1B). DNA sequences are tokenized and embedded with rotary positional encoding and processed by self-attention layers. Protein sequences are embedded with a pre-trained ProteinBERT model (S1A Fig), whereas protein secondary structures are encoded using a dedicated transformer encoder (S1B Fig). Two cross-attention layers are applied to integrate protein sequence and secondary structural information into the DNA sequence representation. The model parameters are optimized to classify protein-DNA pairs as occupied or unoccupied.

We trained COP on 62 mouse C2H2-ZFPs with 2,464 ChIP-seq datasets. For each protein, we curated 3,000 high-confidence occupied sites, yielding a dataset of 186,000 positive (occupied) protein-DNA pairs. To construct negative controls, we combined each DNA sequence with non-occupying proteins, applying gradient-based sampling to select challenging negative pairs with high prediction loss in early epochs, yielding a dataset of 11,346,000 negative (unoccupied) protein-DNA pairs. This strategy generated a total of 11,532,000 protein-DNA pairs for the training set (Fig 1B). Benchmark evaluations demonstrated that COP achieved higher accuracy (63.6%) compared to LightGBM (59.4%) [31], AdaBoost (59.2%) [32], and other known models [24, 33-39] (Fig 1C). These results show that COP outperforms traditional machine learning models in terms of prediction accuracy.

To predict which C2H2-ZFPs may be involved in CTCF-mediated regulation of *cPcdh* gene expression, we applied COP to the 54 pCBS elements of the *cPcdh* promoters. Each pCBS element was fed into COP to predict localized C2H2-ZFPs (Fig 1D). Amino acid sequences and secondary structures were collected for 370 C2H2-ZFPs from UniProt [40] and AlphaFoldDB [41], respectively, and input together with pCBS sequences into COP to infer whether individual C2H2-ZFPs occupy pCBS elements of the *Pcdh* clusters. This revealed 34 C2H2-ZFPs (Figs 1E and S1C-S1L), with 12 trained on COP and 22 untrained (S2 Fig), enriched at the 54 pCBS elements. Among the 34 C2H2-ZFPs, Wiz contains the highest number of 12 ZFs (Figs 1E, 1F, and S1L). Wiz is of particular interest regarding *cPcdh* regulation owing to its high expression levels in the brain [26].

### Wiz colocalizes with CTCF at CBS elements of cPcdh promoters and enhancers

To evaluate occupancy of Wiz at the *cPcdh* gene complex, we generated stable clones with Myc-tagged endogenous *Wiz* via DNA-fragment editing by CRISPR screening 254 single-cell N2a clones (S3A Fig) [13, 42]. We then performed ChIP-seq experiments with a specific antibody against Myc to map Wiz enrichments and found that Wiz is colocalized with CTCF and cohesin at the c*Pcdh* promoter and super-enhancer regions (Fig 2). In the *Pcdhα* cluster, Wiz is enriched at both pCBS and eCBS elements of *Pcdh α1, α8, α9, α12*, and *αc1* as well as two CBS elements (*a* and *b*) flanking the *HS5-1* super-enhancer (Fig 2A). In the *Pcdhβ* cluster, Wiz is enriched at all pCBS elements (Fig 2B). In the *Pcdhγ* cluster, Wiz is enriched at the pCBS elements of *Pcdh γa3, γb1, γb8, γa12*, and *γc3* as well as eight CBS elements of *a-h* within the downstream super-enhancer (Fig 2C).

**Fig 2.**
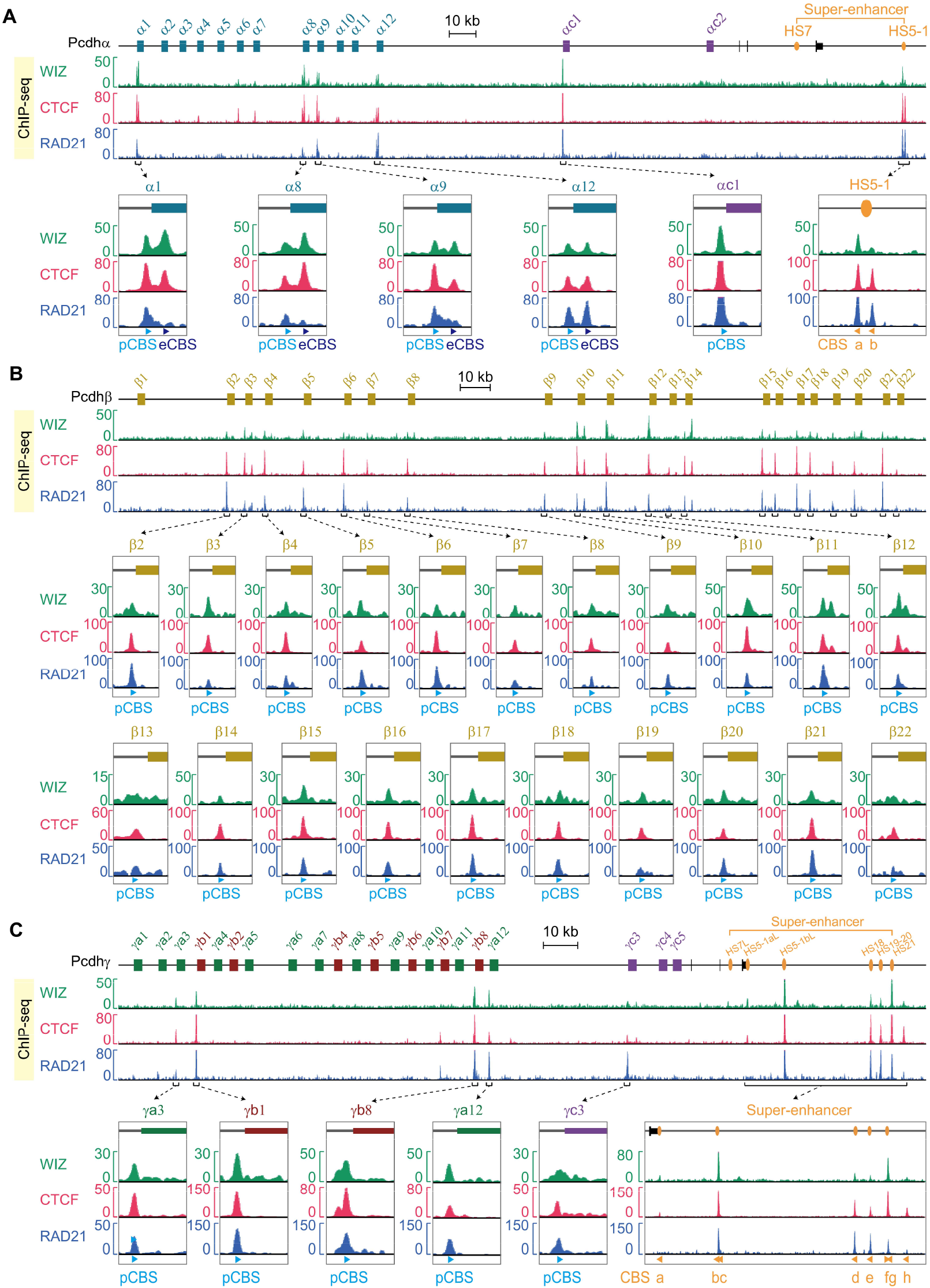
Wiz colocalizes with CTCF and cohesin at both promoter and enhancer regions of the *cPcdh* locus. (**A**-**C**) ChIP-seq profiles of Wiz, CTCF, and RAD21 at the *Pcdh α* (**A**), *β* (**B**), and *γ* (**C**) clusters in N2a single-cell clone expressing endogenous Wiz with a C-terminal Myc tag. The pCBS and eCBS elements within individual genes as well as enhancer CBS elements are highlighted in close-up views.

### Wiz ablation leads to increased cPcdh gene expressions in vitro

We employed DNA fragment editing to delete the entire *Wiz* coding region and obtained two homozygous *Wiz*-knockout (Δ*Wiz*) clones via screening 128 CRISPR single-cell N2a clones (S3B Fig) [13, 42]. We then performed RNA-seq experiments and found 2,377 genes with differential expression patterns (1,831 upregulated and 546 downregulated) (Fig 3A). Remarkably, 1.75% (32 *cPcdh* genes) of the upregulated genes are members of the *Pcdh* clusters, resulting in a unique, ∼32-fold enrichment for upregulated transcripts at this gene complex in comparison with the rest of the genome (Poisson test, *P* = 5.07 × 10^-41^, as determined by a 1-Mb sliding window applied to n = 1,831 transcripts) (Fig 3B). Specifically, members of the *Pcdhα* (Fig 3C), *Pcdhβ* (Fig 3D), and *Pcdhγ* (Fig 3E) clusters are upregulated as shown in heatmap (Fig 3F). In particular, every member of the *Pcdhβ* cluster is upregulated upon *Wiz* deletion (Fig 3D, 3F, and 3G), conspicuously similar to the *Pcdhβ* upregulation upon *Wapl* conditional knockout or knockdown [6, 13]. Finally, ChIP-seq experiments revealed significant increases of active histone marks of H3K4me3 and H3K27ac at the promoter and super-enhancer regions of the *Pcdh* clusters upon *Wiz* deletion (S4 Fig), consistent with *cPcdh* gene upregulation. Collectively, these data suggest that Wiz acts as a transcriptional repressor for the *cPcdh* genes in N2a cells *in vitro*.

**Fig 3.**
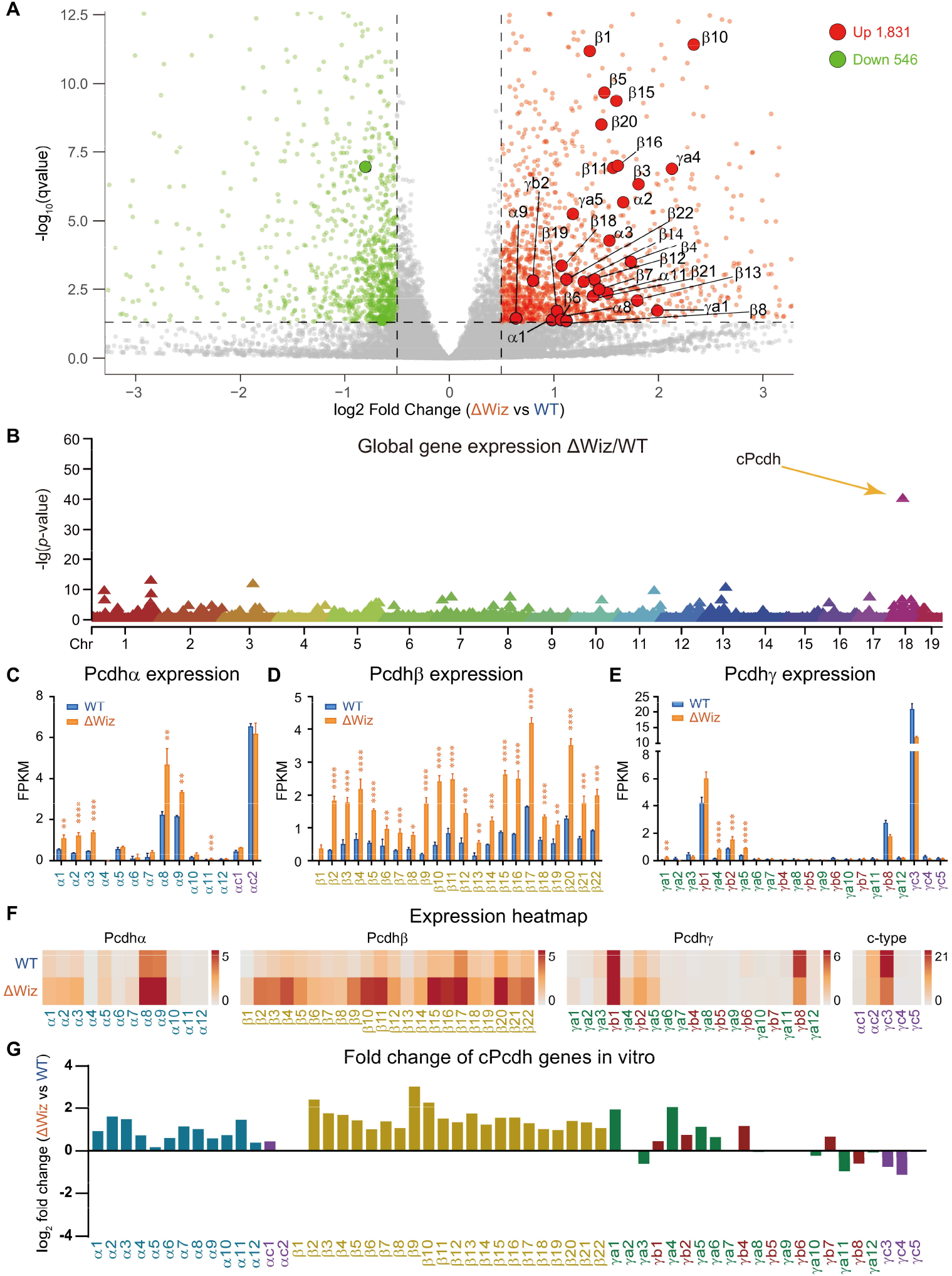
Loss of *Wiz* increases *cPcdh* expression levels in N2a cells *in vitro*. (**A**) Volcano plot depicting differentially expressed genes in neuronal N2a cells upon *Wiz* knockout. Red and green dots represent up-regulated and down-regulated genes, respectively, in Δ*Wiz* single-cell clones compared to the wild-type (WT) control clone. (**B**) Manhattan plot displaying -log_10_ (*q*-values) from the RNA-seq differential expression analysis, showing the strongest enrichment of upregulated genes (1-Mb sliding window) at the *Pcdh* gene clusters in Δ*Wiz* versus WT N2a cell clones. (**C-E**) RNA-seq expression levels of members of the *Pcdh α* (**C**), *β* (**D**), and *γ* (**E**) gene clusters in Δ*Wiz* compared to WT cell clones. (**F**) Heatmaps showing increased expression levels of *cPcdh* upon *Wiz* deletion. (**G**) Bar plots depicting fold changes of gene expression levels of *cPcdh* upon *Wiz* deletion. FPKM, fragments per kilobase of exon per million reads mapped. Data as mean ± standard deviation (SD); unpaired Student’s *t*-test. \**p* ≤ 0.05, \*\**p* ≤ 0.01, ***p ≤ 0.001, \*\*\*\**p* ≤ 0.0001.

### Wiz deletion in the brain leads to increased cPcdh gene expressions in vivo

We next generated *Wiz* conditional knockout mice to investigate its role in the brain *in vivo* because constitutional homozygous null allele of *Wiz* leads to embryonic lethality at early development stages [43]. We first established a *Wiz*-floxed mouse strain (*Wiz*^*+/f*^) using CRISPR/Cas9-mediated homologous recombination via pronuclear microinjection. Briefly, two donor DNA templates were co-injected with Cas9 mRNA and dual single-guide RNAs (sgRNAs) targeting intronic regions flanking the *Wiz* exons 6-13 (S5A Fig). Each donor carried a single *loxP* site, enabling precise insertion of one *loxP* site at flanking intronic regions. Genotyping of 28 P0 chimeric founders identified ten mice carrying correctly targeted *Wiz*-floxed alleles. These chimeric mice were crossed with WT C57BL/6J to establish a germline-transmitted *Wiz*^*+/f*^ line. Specifically, four heterozygous *Wiz*^*+/f*^ mice with two simultaneous *loxP* insertions in a single allele were obtained (S5B Fig) and subsequently crossed with *Emx1*-*Cre* (Empty spiracles homeobox 1-Cre) transgenic mice to generate *Wiz*^*f/f*^;*Emx1*-*Cre* progeny. Further crossing of these mice with *Wiz*^*+/f*^ animals yielded cortex-specific *Wiz* knockouts (*Wiz*^-/-^).

RNA-seq analyses uncovered widespread transcriptional alterations *in vivo* in the cerebral cortices of *Wiz*^-/-^ mice compared with control littermates. At P0, a total of 4,482 genes were differentially expressed upon *Wiz* deletion (2,719 downregulated vs. 1,763 upregulated) (Fig 4A). In particular, 1.53% (27 *cPcdh* genes) of the upregulated genes are mapped within the *cPcdh* locus, far exceeding their expected representation. Specifically, a ∼15-fold enrichment was observed relative to the genomic background (Poisson test, *P* = 1.71 × 10^-25^), based on a 1-Mb sliding-window analysis across the 1,763 upregulated transcripts (Fig 4B). Members of the *Pcdh α* (Fig 4C), *β* (Fig 4D), and *γ* (Fig 4E) clusters are upregulated as shown in heatmap (Fig 4F) and fold changes (Fig 4G). Together, these *in-vitro* and *in-vivo* data clearly demonstrated that *Wiz* represses *cPcdh* gene expression.

**Fig 4.**
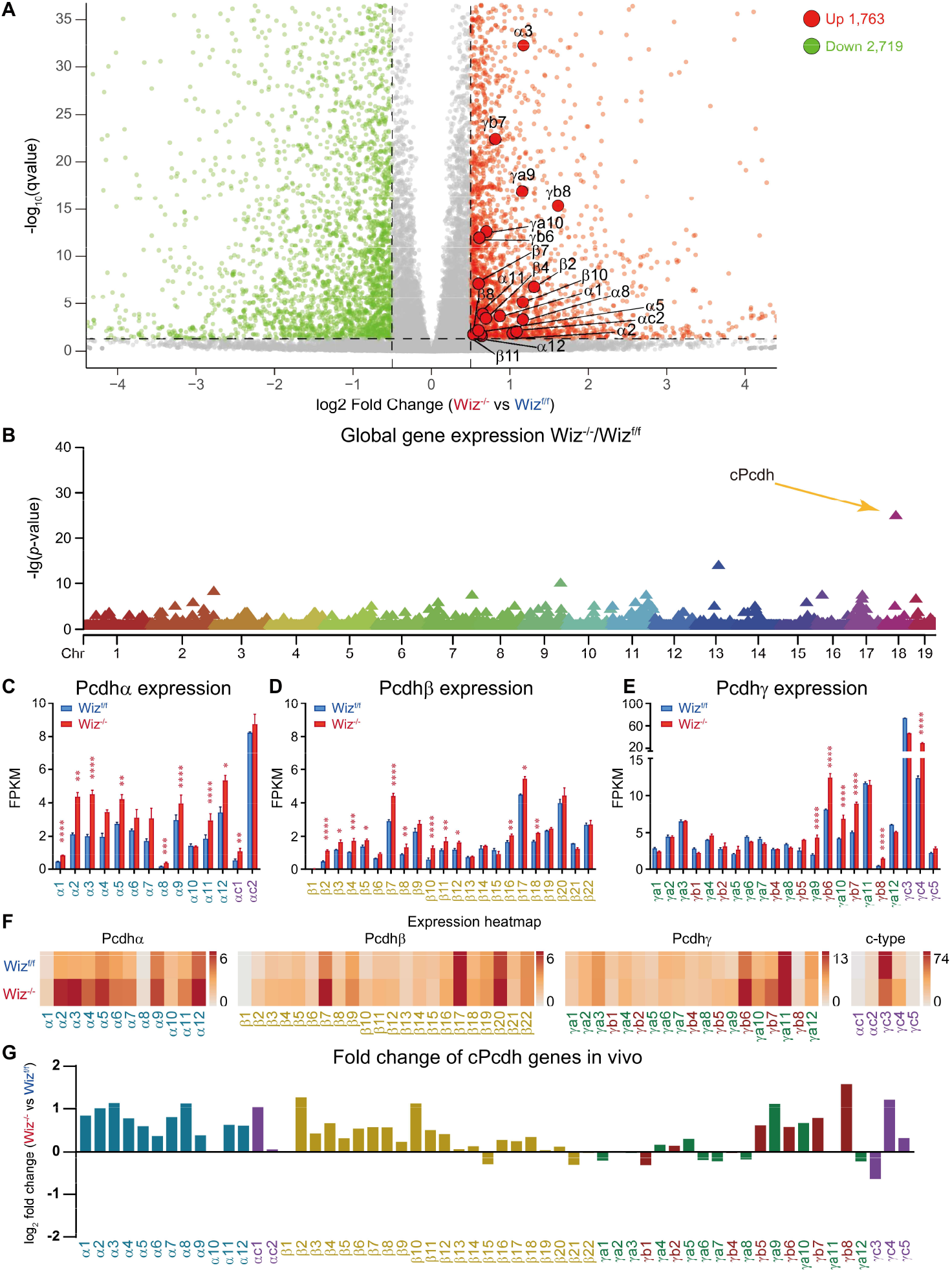
Loss of *Wiz* increases *cPcdh* expression levels in mouse brain *in vivo*. (**A**) Volcano plot depicting differentially expressed genes in the postnatal day 0 (P0) mouse cortices upon conditional knockout of *Wiz*. Red and green dots represent up-regulated and down-regulated genes, respectively, in conditional *Wiz* knockout (*Wiz*^-/-^) versus the *Wiz*^f/f^ control mice. (**B**) Manhattan plot displaying -log_10_ (*q*-values) from the RNA-seq differential expression analysis, showing the strongest enrichment of upregulated genes (1-Mb sliding window) at the c*Pcdh* locus in P0 cortices from *Wiz*^-/-^mice versus *Wiz*^f/f^ controls. (**C-E**) RNA-seq expression levels of members of the *Pcdh α* (**C**), *β* (**D**), and *γ* (**E**) gene clusters in *Wiz*^-/-^mice compared to *Wiz*^f/f^controls, showing a significant increase in c*Pcdh* expression levels upon *Wiz* knockout in the cerebral cortices. (**F**) Heatmaps showing increased *cPcdh* expression levels upon *Wiz* knockout *in vivo*. (**G**) Bar plots depicting fold changes of gene expression levels of *cPcdh* by *Wiz* knockout *in vivo*. FPKM, fragments per kilobase of exon per million reads mapped. Data as mean ± standard deviation (SD); unpaired Student’s *t*-test. \**p* ≤ 0.05, \*\**p* ≤ 0.01, ***p ≤ 0.001, \*\*\*\**p* ≤ 0.0001.

### Wiz represses cPcdh by restricting CTCF/cohesin enrichments at promoters and enhancers

We performed ChIP-seq experiments to assess CTCF/cohesin enrichments upon *Wiz* deletion and found a significant increase in CTCF and Rad21 (a subunit of cohesin) enrichments at the *cPcdh* locus (Fig 5). Specifically, in the *Pcdhα* cluster, CTCF and cohesin enrichments markedly increased at both pCBS and eCBS elements of *Pcdh α1, α8, α9, α12*, and *αc1* as well as at the two CBS elements within the *HS5-1* super-enhancer region (Fig 5A-5C). In addition, there is a remarkable increase of CTCF and cohesin enrichments at the pCBS element of each member of the *Pcdhβ* cluster (Fig 5 D and 5E).

**Fig 5.**
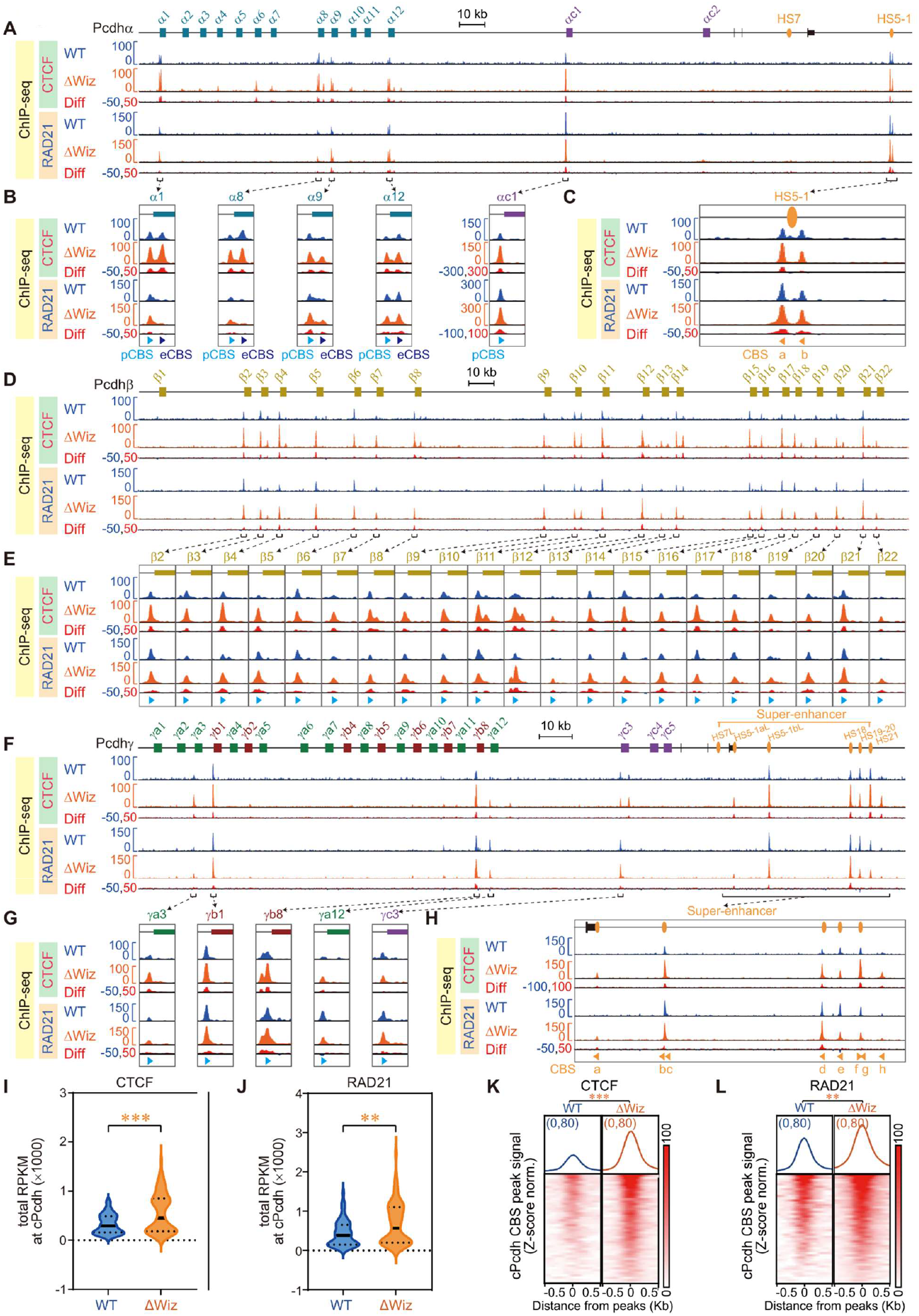
Increased enrichments of CTCF and RAD21 at the *cPcdh* CBS elements upon *Wiz* knockout. (**A**-**H**) ChIP-seq profiles of CTCF and RAD21 at the *Pcdh α* (**A-C**), *β* (**D** and **E**), or *γ* (**F-H**) gene cluster. (**I** and **J**) ChIP-seq quantifications showing a significant increase of CTCF (**I**) and RAD21 (**J**) enrichments upon *Wiz* deletion. (**K** and **L**) Aggregated peak analyses showing a significant increase of CTCF (**K**) and RAD21 (**L**) enrichments at the *cPcdh* locus upon *Wiz* deletion.

Moreover, there is a similar increase throughout the *Pcdhγ* cluster at both variable promoters and downstream super-enhancer (Fig 5F-5H). Finally, total reads and aggregated peak analyses showed a significant increases of both CTCF (Fig 5I and 5K) and cohesin (Fig 5J and 5L) enrichments within the *cPcdh* gene complex upon *Wiz* deletion. In summary, these data suggest that *Wiz* represses *cPcdh* gene expression by restricting CTCF/cohesin enrichments at the CBS elements of promoters and super-enhancers.

### Wiz regulates cPcdh gene expression in a genomic-distance biased manner

Given the emerging role of cohesin-mediated loop extrusion in controlling *cPcdh* transcription via long-range enhancer-promoter communication [4, 6, 9-12], we next examined whether the transcriptional activation of *cPcdh* genes upon *Wiz* loss exhibits positional dependency along the linear genomic sequences. Indeed, analysis of RNA-seq data revealed a pronounced genomic-distance biased increase in *cPcdh* expression levels in both N2a cells *in vitro* (Fig 6A) and mouse cortices *in vivo* (Fig 6B) upon *Wiz* deletion, with enhancer-distal members displaying stronger upregulation than proximal ones. These data suggest that *Wiz* restricts or inhibits *cPcdh* gene expression in a genomic-distance biased manner.

**Fig 6.**
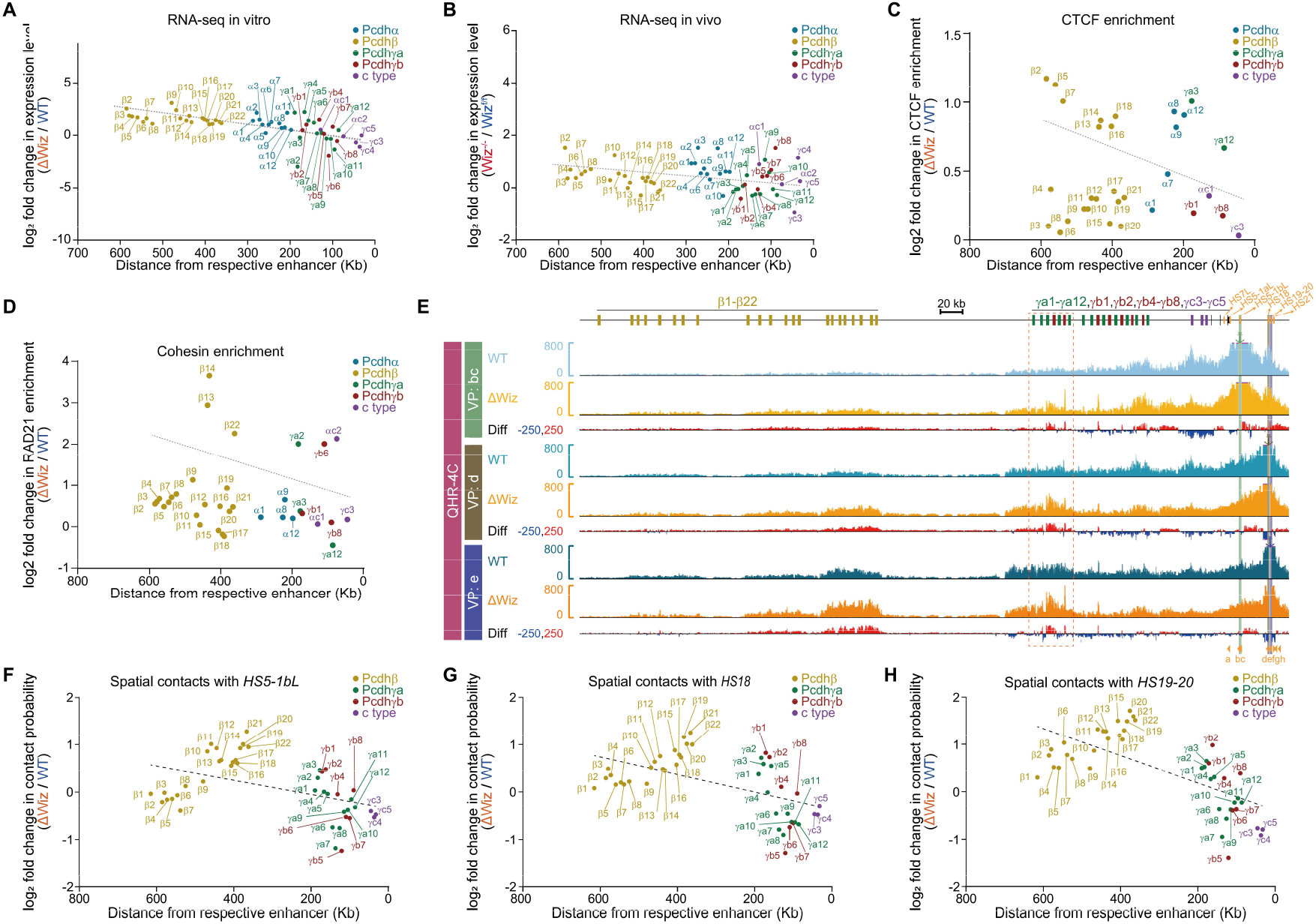
Wiz ablation increases *cPcdh* expression in a genomic-biased manner. (**A**) Scatter plot depicting expression changes of *cPcdh* genes in Δ*Wiz* versus wild-type (WT) cell clones *in vitro*, plotted against their linear genomic distances from respective enhancer. (**B**) Scatter plot depicting expression changes of *cPcdh* genes in P0 cortices from *Wiz*^-/-^versus *Wiz*^f/f^ mice *in vivo*, plotted against their linear genomic distances from respective enhancer. (**C** and **D**) Scatter plots showing genomic distance-biased increase of CTCF (**C**) and RAD21 (**D**) enrichments at pCBS elements of individual c*Pcdh* genes upon *Wiz* deletion. (**E**) 4C profiles using *CBSbc, CBSd*, or *CBSe* as a viewpoint (VP) in Δ*Wiz* compared to WT cells, showing increased chromatin contacts with the distal *Pcdhβγ* genes upon *Wiz* deletion. Differences (Δ*Wiz* versus WT) are shown below the 4C profiles. (**F-H**) Scatter plot showing a genomic-distance biased increase in contact probabilities of *Pcdhβγ* genes with *HS5-1bL* (**F**), *HS18* (**G**), or *HS19-20* (**H**) enhancer within the super-enhancer upon *Wiz* deletion.

To investigate the underlying mechanism, we quantified CTCF and Rad21 ChIP-seq datasets and found a genomic-distance biased increase of CTCF/cohesin occupancy at *cPcdh* genes upon *Wiz* deletion (Fig 6C and 6D), suggesting that *Wiz* may stall cohesin loop extrusion. We then performed a series of QHR-4C (Quantitative high-resolution chromosome conformation capture copy) experiments using the element of CBS*bc*, CBS*d*, or CBS*e* within the *Pcdhβγ* super-enhancer region as a viewpoint and found a significant increase of long-distance chromatin interactions with *Pcdhβ* and distal members of the *Pcdhγ* cluster (Fig 6E). Quantitative 4C analyses revealed a genomic-distance biased increase of spatial contacts between diverse variable promoters and downstream super-enhancers (Fig 6F-6H). Together, these data suggest that *Wiz* restricts or inhibits cohesin loop extrusion from super-enhancer to tandem promoters.

## Discussion

The *cPcdh* gene complex encodes an enormous diversity of cadherin-like cell adhesion proteins functioning as a sophisticated “molecular barcode” system essential for establishing the precise architecture of the mammalian brain [1, 4, 6, 7, 44-47]. These cell-surface adhesion molecules undergo stochastic and combinatorial expression, providing each neuron with a unique tag of biochemical identity for self/non-self discrimination [48-53]. This molecular diversity is also the foundation for neuronal self-avoidance, a process where sister dendritic branches from the same neuron recognize one another via homophilic Pcdh interactions and subsequently repel apposite membranes, ensuring the neuron maximally covers its territory without self-entanglement [54-58]. Beyond self-nonself discrimination and self-avoidance, cPcdhs also mediate synaptic pruning and the refinement of complex neural circuits [5, 59]. When this molecular barcode is disrupted — often due to aberrant 3D genome organization or epigenetic dysregulation — the resulting “miswiring” of the brain can lead to neurodevelopmental disorders such as depression, autism, and schizophrenia [60, 61]. The unique variable and constant genome architecture and great differential distances of tandem variable promoters to the downstream distal super-enhancers impose intrinsic biases on promoter choice and isoform selection [1, 4, 6, 11]. Fine tuning cohesin processivity along the linear chromatin and balanced spatial contacts of variable promoters with distal super-enhancers via topological chromatin insulators are key to overcome the intrinsic genomic-distance biases [7, 12, 13].

Diverse C2H2-type ZFPs constitute the largest family of transcription factors in the mammalian genome [18, 19]. Each ZF consists of a conserved ββα fold stabilized by a single zinc ion coordinated by two cysteines of each β-sheet and two histidines of an α-helix. Although the DNA recognition code is incompletely understood, each ZF domain recognizes a DNA triplet via four specific residues of an α-helix located on the opposite side of the zinc ion. The true power of this family lies in its modular versatility: by linking multiple ZFs in tandem as a daisy chain, these proteins can recognize long, specific DNA elements with diverse affinities, as seen for the 11-finger protein CTCF in 3D genome architecture. We hypothesized that members of the ZFP family may collaborate with CTCF to fine tune the delicate expression patterns of *cPcdh* genes and proposed an AI-based model of COP to identify CTCF-colocalizing C2H2-ZFPs, revealing *Wiz* with the highest number of 12 ZFs. Although Wiz contains 12 C2H2-ZFs, none of them are clustered. Its individual ZFs may contribute to CBS enrichments via protein-protein interactions or RNA binding rather than direct DNA binding. For example, Wiz C-terminal ZF is essential for interacting with the G9a/GLP complex [27, 28]. RNA binding was recently shown to be prevalent among C2H2-ZFPs [62-65]. In a UV crosslinking and immunoprecipitation (CLIP) experiment, 148 of 150 analyzed C2H2-ZFPs were observed to bind directly to RNA in human cells [65].

Wiz was originally identified by a homologous screening [26] and later was shown to maintain H3K9me1/2-marked heterochromatin by stabilizing the G9a/GLP histone methyltransferase complex on chromatin [27, 28]. Wiz-haploinsufficiency results in a reduced level of the *Pcdhβ* gene expression, suggesting that Wiz functions as a transcriptional activator [66]. However, homozygous *Wiz* deletion showed prominent increases of the *Pcdhβ* genes in cultured N2a cells *in vitro* and in mouse brain *in vivo* (Figs 3 and 4). This inconsistency may result from the different sizes of the deleted *Wiz* regions. Our *Wiz* knockout mice targeted the complete set of six C-terminal ZFs (S5B Fig), while previous *Wiz* N-Ethyl-N-nitrosourea (ENU) mutagenesis causes frameshifts of only the two C-terminal ZFs [43, 66].

Finally, integrated and quantitative analyses suggest that Wiz regulates *cPcdh* gene expression by restricting CTCF/cohesin loop extrusion from distal super-enhancers to diverse variable promoters in a genomic-distance biased manner (Figs 5 and 6), explaining the previous observation that distal enhancers preferentially regulate distal *cPcdh* genes whereas proximal enhancers tend to contact with close promoters [12, 67].

Wiz is both a transcription factor and a structural regulator of chromatin loops [28, 30, 66, 68]. Since Wiz colocalizes and interacts with CTCF and cohesin complex [29, 30, 68], Wiz may stabilize G9a/GLP H3K9 methylation enzymes on chromatin regions containing CBS elements via its C-terminal ZF [27, 28, 68] and recruit the CtBP corepressor complex via its three PXDLS-like motifs to silence genes [28]. However, we did not observe obvious alteration of H3K9 methylations at the *cPcdh* locus upon *Wiz* deletion (S6 Fig). Wiz colocalizes with CTCF at insulators, promoters, and enhancers to regulate their spatial interactions [29, 30]. The high conservation of human and mouse Wiz polypeptides (84% identity) and the dispersed organization of 12 ZFs with long linkers is consistent with its architectural or structural role in chromatin looping [25, 30, 66], similar to the structure role at pericentric heterochromatin region of the dispersed ZF protein ZFP512 [69]. Wiz represses *cPcdh* gene expression not by “heterochromatinization” of the locus via H3K9 methylation but by stalling cohesin processivity or loop extrusion, affecting predominantly members of the *Pcdhβ* cluster owing to its great distance from the downstream distal super-enhancer. This could explain the more pronounced increases of spatial contacts of super-enhancers with distal promoters than with proximal promoters, providing a mechanistic explanation for the preferential activation of distal *cPcdh* genes upon *Wiz* deletion.

Collectively, these observations point to a previously unrecognized architectural role of Wiz in modulating chromatin topology. By restricting longer-range enhancer-promoter contacts across the three *Pcdh* clusters via constraining cohesin chromatin loop extrusion, Wiz regulates *cPcdh* expression in a genomic-distance biased manner.

## Materials and methods

### Animals

All mouse strains were maintained at 23 °C on a 12/12 h light-dark cycle (7:00-19:00) in an SPF (specific pathogen free) facility. All experiments were approved by the Institutional Animal Care and Use Committee (IACUC) of Shanghai Jiao Tong University (protocol#: 1602029).

### Plasmid construction

Plasmids used for expressing sgRNA were constructed as previously described [42, 70]. In brief, each pair of complementary oligonucleotides (S1 Table) was annealed in NEBuffer 2 (NEB, B7002S) to produce a double-stranded DNA (dsDNA) fragment with 5’ overhangs of ‘ACCG’ and ‘AAAC’ at two ends, respectively. Each annealed dsDNA fragment was cloned into the *Bsa*I site of pGL3 vector under the U6 promoter.

For generation of single-cell clones with *Wiz* endogenously Myc-tagged at the C-terminus, a donor plasmid was designed for Cas9-mediated homologous recombination. Left and right homologous arms were amplified from the mouse genomic DNA by PCR using a pair of primers (S1 Table) and cloned into the *Eco*RI and *Xba*I sites of the pTNT vector (Promega, L5610). The Myc-tag coding sequence was inserted between the left and right homologous arms by overlapping PCR using primers containing the Myc-tag coding sequence (S1 Table).

For generation of *Wiz*-floxed mice, two donor plasmids were designed for targeting the fourth and the tenth introns of *Wiz*, respectively. For each donor, the left and right arms were amplified by PCR from the mouse genomic DNA, respectively, with the reverse primer of the left arm and the forward primer of the right arm containing complementary *loxP* sequences. The amplified products of left and right homology arms were assembled into the linearized pUC19 vector using the Clone Express MultiS OneStep Cloning Kit, to generate the final donor plasmid with the *loxP* sequence flanked by the two homologous arms. All plasmids were confirmed by Sanger sequencing.

### Cell culture

Mouse Neuro-2a (N2a) cells were cultured in modified Eagle’s medium (Gibco, 11095080) containing 10% fetal bovine serum (Sigma-Aldrich, F0193), 1% penicillin–streptomycin (Gibco, 15140122), and 1% non-essential amino acids (Gibco, 11140050). Cells were maintained at 37 °C in a humidified environment with 5% CO_2_.

Cells were passaged when reaching approximately 80% confluence. After removal of the medium, cells were gently washed once with 1 ml of 1× PBS (Gibco, 70011044). Detachment was achieved by adding 1 ml of 0.25% trypsin–EDTA (Gibco, 25200056) and incubating the cells at 37 °C for 3 min. Enzymatic digestion was terminated by the addition of 1 ml of complete medium. The cell suspension was transferred to a 15-ml conical tube and centrifuged at 900 rpm for 5 min at room temperature. Following centrifugation, the supernatant was removed and the cells were resuspended in 1 ml of fresh medium. For subculturing, 250 µl of the resulting cell suspension was seeded into a new dish containing 5 ml of complete medium.

### Generation of single-cell clone with Wiz endogenously Myc-tagged

To tag Myc at the C-terminus of endogenous Wiz, sgRNA and donor plasmids were co-transfected with the Cas9-expression plasmid into the N2a cells, using Lipofectamine 3000 (Invitrogen, L3000015) according to the manufacturer’s protocol. Transfected cells were cultured for 2 days, followed by selection with 2 μg/ml puromycin for four days. Cells were subsequently transferred to the puromycin-free medium and maintained for one week for recovery. After recovery, the transfected population cells were genotyped by PCR to first confirm the CRISPR insertion. Specifically, genomic DNA was extracted and used as a template for PCR amplification with specific primers (S1 Table) to detect the targeted insertion of Myc-tag sequence. Cell populations with targeted insertion were then subjected to single-cell clone screening. Briefly, the transfected cells were diluted and seeded into the 96-well plates at approximately one cell per well. A total of 254 single-cell clones were screened, and one homozygous single-cell clone was obtained for Myc-tagging at the C-terminus of endogenous Wiz. All single-cell clones were genotyped by Sanger sequencing.

### Generation of ΔWiz single-cell clones

To knock out *Wiz*, a pair of sgRNAs was designed to target the first and the tenth introns of *Wiz* that span the coding sequence of ZF1-12. The two sgRNA-expression plasmids were co-transfected with the Cas9-expression plasmid into the N2a cells, using Lipofectamine 3000. Transfected cells were cultured, puromycin-selected, and subjected to single-cell clone screening as described above. Genomic DNA was extracted and PCR-amplified using specific primers (S1 Table) to detect the targeted deletion of *Wiz*. A total of 128 clones were screened and two homozygous Δ*Wiz* single-cell clones were obtained. All clones were genotyped by Sanger sequencing.

### Synthesis of sgRNA and Cas9 mRNA for microinjection

Both sgRNA and Cas9 mRNA were synthesized via *in-vitro* transcription. For sgRNA transcription, a DNA template containing a T7 promoter followed by sgRNA targeting sequence and scaffold sequence was PCR-amplified from the pGL3 plasmid with a pair of primers (S1 Table). The amplified template was purified with the QIAquick PCR Purification kit (Qiagen, 28104), extracted with phenol-chloroform, precipitated with ethanol, and resuspended in RNase-free water. The sgRNA was transcribed using the MEGAshortscript T7 kit (Invitrogen, AM1354) and purified with the MEGAclear kit (Invitrogen, AM1908) according to the manufacturer’s protocol. The transcribed sgRNA was eluted in 30 μl of Elution buffer, quantified with a NanoDrop 2000 Spectrophotometer, and aliquoted at 2.5 μg per tube for storage at −80 °C to avoid repeated freeze-thaw cycles.

For Cas9 mRNA *in-vitro* synthesis, the pcDNA3.1-Cas9 vector, containing Cas9 coding sequence under a T7 promoter, was linearized with *Xba*I (NEB, R0145S), purified by gel extraction, extracted with phenol-chloroform, and precipitated by ethanol. After ethanol precipitation, the plasmid DNA pellet was resuspended in 15 μl of RNase-free water and used for *in-vitro* transcription with the mMACHINE T7 ULTRA kit (Invitrogen, AM1345). The transcribed Cas9 mRNA was then subjected to DNase digestion, poly(A) tailing, and purification with the MEGAclear kit. Purified Cas9 mRNA was quantified by NanoDrop 2000 Spectrophotometer and aliquoted at 5 μg per tube for storage at −80 °C.

### Generation of Wiz-floxed mice for conditional knockout of Wiz

C57BL/6J female mice of 3-6 weeks old were induced for superovulation by subjecting to an intraperitoneal injection of 10 U of pregnant mare serum gonadotropin (PMSG) (Solarbio, P9970) and 48h-later, a second injection of 10 U of human chorionic gonadotropin (hCG) (MCE, HY-107953). After the injection of hCG, each female mouse was placed in an independent cage with a stud male of C57/BL6J for crossing. Plugged females were euthanized, and fertilized zygotes were collected from oviducts and cultured in M2 medium at 37 °C with 5% CO_2_ for microinjection.

To construct the floxed *Wiz* allele, two sgRNAs were designed to target the introns 5 and 13 of *Wiz*, respectively, to induce Cas9-mediated homologous recombination. Briefly, 1.25 μg each of the two sgRNAs were mixed with 5 μg each of two donor plasmids and 5 μg of Cas9 mRNA in RNase-free water to prepare a 50-µl injection mixture. For microinjection, chambers containing M2 medium drops equilibrated under mineral oil were prepared in advance. Approximately 2 picolitre of the injection mixture was injected into the pronucleus of each zygote. After recovery for 30 min, morphologically normal zygotes were transferred into the oviducts of pseudopregnant ICR females.

### Mouse genotyping

The chimeric F0 mice were genotyped by PCR using specific primers (S1 Table). In brief, a small piece of the mouse tail was snipped into an Eppendorf tube. The tail tissue was lysed in 30 μl of the Solution A (25 mM NaOH) at 95 °C for 20 min and neutralized with 30 μl of the Solution B (25 mM Tris-HCl pH 6.8). The 2 μl of the neutralized tail lysis solution was used as a template to screen for the targeted mutations by polymerase chain reaction (PCR under the conditions: 95 °C, 3 min; 95 °C, 15 s, 58 °C, 15 s, 72 °C, 15 s for 40 cycles; and a final extension at 72 °C, 5 min) using specific primers. PCR products were Sanger-sequenced for genotyping. The chimeric F0 mice with desired insertions were crossed with wildtype C57BL/6J mice to generate heterozygous (*Wiz*^*f/+*^) F1 mice. The targeted F1 male and female mice were crossed to generate the F2 mice. The homozygous (*Wiz*^*f/f*^) F2 mice were genotyped by Sanger sequencing and used for generation of *Wiz* conditional knockout.

### RNA-seq

RNA-seq experiments were performed as previously described with minor modifications [11]. Briefly, cultured cells or dissociated cortical cells were lysed in 1 ml of TRIzol (Invitrogen, 15596026) by vortexing vigorously for 15 min at room temperature. The sample was then centrifuged at 12,000 g for 10 min at 4 °C. The supernatant was transferred to a new microcentrifuge tube, and 0.2 ml of chloroform was added. After vigorous shaking for 15 s, the sample was incubated for 3 min at room temperature. After centrifugation at 12,000 g for 15 min at 4 °C, the aqueous phase was carefully transferred to a new RNase-free tube and mixed with 0.5 ml of isopropanol to precipitate RNA. After a 10-min incubation at room temperature, RNA was precipitated by centrifugation at 12,000 g for 10 min at 4 °C. The precipitated RNA was washed once with 1 ml of 75% ethanol, air-dried for 5 min, and resuspended in 30 μl of nuclease-free water. The concentration of RNA was measured using a spectrophotometer (NanoDrop, 2000). The high-quality RNA, with an A260/A280 ratio of ∼2.0, was subjected to library preparation.

RNA-seq libraries were prepared using the Universal V6 RNA-seq Library Prep kit for Illumina (Vazyme, NR604-01) following the manufacturer’s instructions. Briefly, the polyadenylated mRNA was isolated using oligo(dT) coupled to magnetic beads (Vazyme, N401) from 100 ng of total RNA and fragmented by heating at 94 °C for 8 min. The first cDNA strand was synthesized by reverse transcription with a random primer (N6). After synthesizing the second strand of DNA, the adapter was ligated. The ligated DNA was cleaned by AMPure XP Beads (Beckman, A63881), and mixed with 5 μl of the P5 primer, 5 μl of the P7 primer, and 25 μl of HiFi Amplification Mix (Vazyme, NR604) for PCR-amplification (98 °C, 30 s; 98 °C, 10 s, 60 °C, 30 s, 72 °C, 30 s for 10 cycles; and a final extension at 72 °C, 5 min). RNA-seq library was sequenced on an Illumina platform.

### ChIP-seq

Cultured cells were first washed twice with PBS, detached by trypsin digestion, resuspended in 10 ml of medium to neutralize the trypsin digestion. Formaldehyde (Thermo, 28908) was added to a final concentration of 1% for cross-linking at room temperature for 10 min. The glycine was added to a final concentration of 125 mM and incubated at room temperature for 5 min to quench the cross-linking reactions. Cross-linked cells were centrifuged at 2,500 g for 10 min at 4 °C and washed with ice-cold PBS. Washed cells were lysed on ice using 1 ml of ChIP Lysis Buffer (10 mM Tris-HCl pH 7.5, 1 mM EDTA, 1% Triton X-100, 0.1% sodium deoxycholate, 150 mM NaCl, 1× protease inhibitors) for 30 min, and centrifuged at 2,500 g for 5 min at 4 °C to obtain cell nuclei.

The isolated nuclei were resuspended in 0.7 ml of ChIP Lysis Buffer and sonicated using a Bioruptor Plus sonicator (Diagenode) in a non-contact mode at high power at a train of 30 cycles of 30 s ON/30 s OFF to yield 200-1,000 bp DNA fragments. The sonicated samples were centrifuged at 14,000 g for 10 min at 4 °C. The supernatants were transferred to a new tube and precleared with 50 μl of protein A/G-agarose beads (Millipore, 16-157) at 4 °C with slow rotation for 2 h. After centrifugation at 2,000 g for 1 min at 4 °C, the protein A/G-agarose beads were discarded, and the supernatants were transferred to a new tube. The primary antibody was added and incubated overnight at 4 °C with slow rotation for immunoprecipitation. 50 μl of the protein A/G-agarose beads were added and incubated at 4 °C with slow rotation for 3 h. The samples were centrifuged at 2,000 g for 1 min at 4 °C and followed by sequential washes with Low Salt Washing Buffer (0.1% SDS, 1% Triton X-100, 2 mM EDTA, 20 mM Tris-HCl pH 8.0, 150 mM NaCl), High Salt Washing Buffer (0.1% SDS, 1% Triton X-100, 2 mM EDTA, 20 mM Tris-HCl pH 8.0, 500 mM NaCl), LiCl Washing Buffer (0.25 M LiCl, 1% NP-40, 1% sodium deoxycholate, 1 mM EDTA, 10 mM Tris-HCl pH 8.0), and TE Buffer (10 mM Tris pH 8.0, 1 mM EDTA).

The washed antibody/protein/DNA complexes were eluted twice with 100 μl of Elution Buffer (50 mM Tris-HCl pH 8.0, 10 mM EDTA, 1% SDS) by incubation at 65 °C for 30 min with vortexes. The 200-μl eluted solution was mixed with 200 μl of TE buffer, de-cross-linked at 65 °C overnight with vortexes, and sequentially digested with 2 μl of 10 mg/ml RNase A at 37 °C for 2 h and 8 μl of 10 mg/ml proteinase K at 55 °C for 2 h. The DNA was purified with 400 μl of phenol/chloroform, precipitated, and resuspended in 20 μl of nuclease-free water. DNA concentration was measured by PicoGreen reagents. A total of 10 ng of DNA was used as a template for library construction using NGS Ultima Pro DNA Library Prep Kit (Yeasen, 12201ES96). The generated ChIP-seq library was sequenced on an Illumina platform.

### QHR-4C

QHR-4C experiments were carried out as previously described with minor modifications [12]. Briefly, about 1 × 10^7^ cells were harvested as described above and crosslinked with formaldehyde at a final concentration of 2% at room temperature for 10 min. Crosslinking was quenched by adding 2 M glycine to a final concentration of 200 mM. Crosslinked cells were centrifuged at 220 g for 5 min, washed with 10 ml of ice-cold PBS, and centrifuged again at 220 g for 5 min. Cells were then permeabilized twice with 200 μl of ice-cold 4C permeabilization buffer (50 mM Tris-HCl pH 7.5, 150 mM NaCl, 5 mM EDTA, 0.5% NP-40, 1% Triton X-100, and 1× protease inhibitors) each for 10 min. After centrifugation, the pellet was resuspended in 73 μl of water, 10 μl of 10× *Dpn*II buffer, and 2.5 μl of 10% SDS. The resuspended cells were incubated at 37 °C for 1 h with shaking at 900 rpm. 12.5 μl of 20% Triton X-100 was added into the reaction to quench SDS and incubated at 37 °C for 1 h with shaking at 900 rpm. The cells were then digested *in situ* overnight at 37 °C with 2 μl of 10 U/μl *Dpn* II while shaking at 900 rpm.

After the inactivation of *Dpn* II at 65 °C for 20 min, the pellets of the nuclei were collected by centrifuging at 1,000 g for 1 min, and the supernatant was removed completely, which ensures that the subsequent ligation reaction can be performed in a small volume. Proximity ligation was carried out for 24 h at 16 °C with 1 μl of T4 DNA ligase (400 U/μl) in 100 μl of 1× T4 ligation buffer. The ligated product was then de-cross-linked by heating at 65 °C for 4 h in the presence of 1 μl of proteinase K (10 mg/ml) to digest proteins. The DNA was then extracted using phenol-chloroform. One μl of glycogen (20 mg/ml) was added to facilitate DNA precipitation. The precipitated DNA was resuspended in 50 μl of water and fragmented to an average size of 200-500 bp using a Bioruptor sonicator (low-power setting, 30 s ON and 30 s OFF, 12 cycles).

The fragmented DNA was used as the template for linear amplification using a 5’ biotinylated primer (S1 Table) complementary to the viewpoint fragment. The amplification was performed in a 100-μl PCR reaction (95 °C, 2 min; 95 °C, 15 s, 58 °C, 25 s, 72 °C, 90 s for 82 cycles; and a final extension at 72 °C, 5 min). The PCR products were denatured at 95 °C for 5 min and immediately chilled on ice to generate single-stranded DNA (ssDNA). The generated ssDNA was then enriched and purified with Streptavidin Magnetic Beads (Thermo), followed by ligation with annealed adaptors in a 45-μl ligation reaction (20 μl of DNA-coated beads, 4.5 μl of 10× T4 ligation buffer, 10 μl of 30% PEG 8000, 1 μl of 50 μM adaptor, 0.9 μl of T4 DNA ligase, 8.6 μl of water). After ligation, the beads were washed twice with 1× Binding and Washing Buffer (5 mM Tris-HCl pH 7.5, 1 M NaCl, 0.5 mM EDTA) to remove unligated adaptors, and finally resuspended in 10 μl of water. Using the bead-bound DNA as template, the 4C library was PCR-amplified using high-fidelity DNA polymerase (Vazyme, P505-d1) with specific primers (S1 Table) under the cycling condition (95 °C, 3 min; 95 °C, 15 s, 60 °C, 30 s, 72 °C, 1 min for 19 cycles; and a final extension at 72 °C, 5 min). The amplified 4C library was purified with High Pure PCR Product Purification Kit (Roche) and sequenced on an Illumina platform.

### Artificial intelligence (AI) model Processing of mouse C2H2-ZFP data

All amino acid sequences of mouse C2H2-ZFPs were retrieved from UniProt [40] using the query “(ft_zn_fing: C2H2) AND (organism_id: 10090)”, yielding 380 entries. Predicted 3D protein structure models were obtained from the AlphaFold Protein Structure Database (AlphaFoldDB) [41]. Of these 380 proteins, six lacked corresponding structural models in AlphaFold DB, and four exhibited discrepancies in sequence length between UniProt and AlphaFold DB entries. These ten proteins therefore excluded. Consequently, high-confidence structural and sequence data for 370 mouse C2H2 ZFPs (S2 Table) were retained for downstream analyses.

The 3D protein structures were converted into secondary structure assignments using DSSP [71], which classifies residues into nine canonical secondary structure types: *α*-helix, residue in isolated *β*-bridge, extended strand participating in *β-*ladder, 310-helix, *π*-helix, *κ*-helix (poly-proline II helix), hydrogen-bonded turn, bend, and none (unstructured). Two functionally important motifs, namely, C2H2-ZF motif, essential for sequence-specific DNA binding, and the KRAB domain, commonly found adjacent to C2H2-ZF arrays, were explicitly annotated as distinct secondary structure classes to preserve their biological relevance in downstream analyses.

### Processing of public ChIP-seq data

Among the 370 mouse C2H2-ZFPs, ChIP-seq data were available for 62 in the NCBI SRA (S2 Table). Raw sequencing reads were downloaded and aligned to the mm9 reference genome using Bowtie2 (v2.3.4.1). Peak calling was performed with MACS (v1.4.2), and peaks overlapping with the ENCODE mm9 blacklist [72] were excluded. To consolidate nearby binding sites, peaks with at least 50-bp of overlap were clustered. within each cluster, a single representative peak was retained, especially, the one located at the 90th percentile of *p*-values, to minimize redundancy and suppress spurious peak calling. To ensure uniformity in peak width, the top 10% widest and narrowest peaks were discarded. To further reduce false positives, the top 10% of peaks with the highest *p*-values (least significant) and lowest *p*-values (potentially artifactual) were also excluded. Finally, to balance statistical robustness and computational efficiency, 3,000 peaks were randomly sampled per protein, and 100-bp sequences centered on each peak summit were extracted for downstream analyses.

### Tokenization of input sequences

Input DNA sequences were tokenized using a vocabulary comprising the characters: m, c, A, C, G, T, and N, corresponding to the indices 0-6, respectively. Specifically, m denotes a padded token used for sequence length masking. The character c serves as a sequence classification token, positioned at the beginning of each sequence, and functions as the input to the classification head for downstream prediction, following the BERT-style [CLS] token paradigm [73]. A, C, G, and T represent the four canonical DNA nucleotides. N represents an ambiguous or unknown nucleotide. For each pCBS element within the *cPcdh* genes, the 38-bp core motif was extended symmetrically to a total length of 256 bp prior to input into COP. This extension was guided by transcription factor binding peak data from the ReMap database [74], aiming to maximize coverage of bound C2H2-ZFPs.

Input protein sequences were encoded using ProteinBERT [75] with a vocabulary of 26 tokens (indexed 0-25). To ensure uniform sequence length, a padding token p was introduced. Sequence boundaries were explicitly marked by adding a start token s at the beginning and an end token e at the terminus. The tokens A, C, D, E, F, G, H, I, K, L, M, N, P, Q, R, S, T, V, W, and Y represent the 20 standard amino acids. The remaining tokens, U, X, and o, correspond to selenocysteine, undefined amino acids, and non-canonical amino acids, respectively.

The DSSP-predicted secondary structures of proteins were encoded with a vocabulary of 12 tokens: m, H, B, E, G, I, P, T, S, K, Z, and -, corresponding to indices 0-11, respectively. The token m served as a mask (padding) token. The structural tokens of H, B, E, G, I, P, T, and S correspond to *α*-helix (H), residue in isolated *β*-bridge (B), extended strand participating in *β-*ladder (E), 310-helix (G), *π*-helix (I), *κ*-helix (P), hydrogen-bonded turn (T), bend (S), and none (-), respectively. The remaining tokens, K and Z, denote KRAB domain (K) and C2H2 ZF motif (Z), respectively. This encoding was processed by a transformer encoder module.

### ProteinBERT architecture

The ProteinBERT architecture (S1A Fig) comprises alternating global and local processing modules. The global module captures sequence-level contextual information, whereas the local module generates residue-specific embeddings. Cross-attention mechanisms facilitate bidirectional information exchange between these two modules, maintaining linear computational complexity with respect to sequence length, thereby ensuring scalability and efficiency for long protein sequences. The maximum protein length was set to 2,700 residues. Finally, a projection header was incorporated to align the embedding dimensionality with that required by downstream tasks.

### Protein secondary structure encoder

Protein secondary structural features (S1B Fig) were encoded using a transformer encoder applied iteratively. The resulting outputs were normalized via root-mean-square (RMS) normalization and subsequently served as the key and value embeddings in cross-attention with DNA embeddings.

### COP architecture

COP comprises five modules: a self-attention module for processing DNA embeddings, a cross-attention module that integrates DNA and protein embeddings, a cross-attention module that fuses DNA and secondary structure embeddings, a feedforward network (FFN), and a final classifier.

Input DNA sequences were first tokenized and projected into a 128-dimensional embedding space. These embeddings were then subjected to random dropout, followed by RMS normalization approach used in DeepSeek [76]. The normalized embeddings served as input to a self-attention module equipped with residual connections, enabling the model to capture long-range dependencies and global nucleotide interactions along the DNA sequence. Multi-head attention was implemented using torchtune [77], with rotary positional embeddings (RoPE) applied exclusively to the query and key projections, but not the value embeddings. Additionally, the torchtune implementation includes a linear output projection layer to transform the attention outputs into the desired feature dimension.

The output embedding of the DNA classification token was then RMS-normalized and passed to the downstream cross-attention module. Meanwhile, protein embeddings, extracted from the pretrained ProteinBERT model, were projected into key and value representations, allowing the model to capture contextual interactions between DNA and protein sequences. The resulting DNA-protein cross-attention output was subsequently RMS-normalized and fed into a second cross-attention layer. In this layer, embeddings encoding protein secondary structural features, including C2H2-ZF motifs, served as the key and value inputs, enabling the model to learn associations between protein secondary structure and DNA sequence.

The outputs underwent further processing by an FFN, which comprises an RMSNorm layer, followed by two linear transformations interleaved with a Gaussian error linear unit (GELU) activation function, and concluded with a dropout layer applying a stochastic mask. The entire pipeline, comprising self-attention, cross-attention, and FFN layers, was repeated multiple times. Finally, the embedding corresponding to the DNA classification token was normalized using RMSNorm and fed into the classifier head to produce the final logits.

This architecture encodes DNA sequence, protein sequence, and protein secondary structure representations independently in early stages, while enabling targeted, hierarchical information integration into the DNA representation via cascaded cross-attention. Transfer learning, leveraging the pretrained ProteinBERT weights, consistently yielded superior performance compared to de novo training.

### Positive and negative samples

For each of the 62 C2H2-ZFPs, 3,000 ChIP-seq binding sites were randomly sampled, yielding a total of 62 × 3,000 positive protein-DNA pairs. To generate negative pairs, each protein was paired with binding sites identified for the other 61 proteins, yielding 62 × 61 × 3,000 negative protein-DNA pairs. To address class imbalance, we employed hard negative sampling. Specifically, for each binding site in a batch of *B* positive pairs, the corresponding *B* × 61 negative pairs were filtered to exclude any that fell within a 300 bp window of the binding site. The remaining negatives were then ranked by their loss values accumulated over previous training epochs. The top *R* (50%) highest-loss instances were selected as hard negatives, while the rest were sampled uniformly without replacement. This gradient-based one-sided sampling (GOSS) strategy [31] effectively preserved the overall negative distribution while adaptively enriching the training set with informative, challenging examples.

### Benchmarking

To evaluate the modeling performance (S3 Table), comparisons were made against both classical machine learning algorithms and a previously proposed deep learning method. For classical baselines, LightGBM and several standard classifiers, including support vector machine (with stochastic gradient descent learning), perceptron, passive-aggressive classifier, decision tree, random forest, AdaBoost, and naive Bayes, were implemented. To accommodate memory constraints, negative samples were downsampled to match the number of positive samples. For all baseline models, tokenized DNA, protein, and secondary structure sequences were concatenated as input features.

Ensemble-based methods, LightGBM and AdaBoost (boosting) as well as Random Forest (bagging), achieved the highest accuracies, approximately 0.59. Notably, naive Bayes outperformed linear models of SVM, perceptron, and passive-aggressive classifier, suggesting that C2H2-ZFP-DNA recognition has inherent non-linear dependencies. The deep learning model DeepZF [24] was also benchmarked. DeepZF comprises two components. One is classification of individual C2H2 ZF into DNA-binding versus non-binding categories. The other is prediction of triple-nucleotide motif for a C2H2 ZF. For each protein, the predicted triple-nucleotide motifs from tandem C2H2 fingers were concatenated to generate a composite DNA binding motif. These composite motifs were then scored against positive and negative DNA binding sites using FIMO (Find Individual Motif Occurrences). A score threshold was optimized on 90% of the ChIP-seq dataset and evaluated on the remaining 10%. As shown in S4 Table, DeepZF did not outperform the ensemble learning methods or other common classifiers in predicting binding sites.

### Bioinformatics analyses

#### Data analyses of RNA-seq

RNA-seq raw FASTQ files were aligned to the mouse reference genome (GRCm38) using Hisat2, generating Sequence Alignment Map (SAM) files. These SAM files were converted into sorted, indexed Binary Alignment Map (BAM) files using Samtools (v1.15.1). The generated BAM files were processed using Cufflinks to quantify transcript expression levels in units of fragments per kilobase of variable exon per million fragments mapped (FPKM). Raw read counts were subsequently imported into DESeq2 for differential gene expression analyses, with significantly differentially expressed genes defined as those exhibiting an absolute log_2_ fold change > 0.5 and an adjusted *p*-value < 0.05.

Manhattan plots depicting localized enrichments of downregulated genes were generated using the CMplot package (v4.5.1) in R. The mouse reference genome was partitioned into non-overlapping 1-Mb bins. The genomic locations of downregulated genes were intersected with these bins to calculate the frequency of occurrences within each bin. The distribution of occurrences was modeled using a Poisson distribution, with the maximum-likelihood estimator of λ derived from the mean frequency of events. Probabilities of observing the signals in each bin were then computed under the Poisson model and visualized with CMplot to produce Manhattan plots.

#### Data analyses of ChIP-seq

Raw reads were trimmed using fastp (v0.21.0) to remove the first 10 bp barcode and adaptor sequences. Trimmed reads were then aligned to the mouse reference genome (mm9) using Bowtie2 (v2.3.4.1), generating SAM files. These SAM files were converted, sorted, and indexed into BAM files using Samtools (v1.15.1). For comparative analysis of protein-DNA occupancy across samples, BAM files were normalized to reads per kilobase per million mapped reads (RPKM) with a bin size of 20 bp using the bamCoverage module in deepTools. The resulting bedGraph files were uploaded to the UCSC Genome Browser for visualization of genomic regions of interest. Narrow peaks were identified using MACS (v1.4.2 20120305) with a stringent q-value cutoff of 0.001. Heatmaps depicting signal distribution were generated using the plotHeatmap module of deepTools. To assess enrichment at each *cPcdh* gene, the peak summit within the corresponding pCBS element was identified, and its signal intensity was used as the enrichment score.

#### Data analyses of QHR-4C

For the QHR-4C data, P7 reads were demultiplexed based on unique barcode-index combinations. Anchor primer sequences were trimmed, and PCR duplicates were removed using FastUniq (v1.1). The resulting non-redundant reads were aligned to the mouse reference genome (mm9) using Bowtie2 (v2.3.5) to generate SAM files, which were then converted to sorted, indexed BAM files. BAM files were processed with the r3Cseq package (v1.20) in R (v3.3.3) to compute normalized read counts as reads per million (RPM). BedGraph files generated by r3Cseq were uploaded to the UCSC Genome Browser for visualization. Loop intensities were quantified using the RPKM.

#### Quantification and statistics

RNA-seq, ChIP-seq, and QHR-4C experiments were performed with at least two biological replicates. All statistical tests were calculated using GraphPad (v8.4.2), R (v4.3.3) or Python (v3.6.9) scripts. Data were presented as mean ± standard deviation (SD). The values of statistical significance were calculated using an unpaired Student’s t-test, with significance levels denoted as follows: ‘*’ for p ≤ 0.05, ‘**’ for p ≤ 0.01, ‘***’ for p ≤ 0.001, and ‘****’ for p ≤ 0.0001.

## Acknowledgments

We are grateful to Dr. Yan Jiang for kind gift of Emx1-Cre mice. This work was supported by grants from the National Natural Science Foundation of China (32330016) and the National Key R&D Program of China (2022YFC3400204) to Q.W.

## Author contributions

**Conceptualization:** Qiang Wu.

**Data Curation:** Tianjie Li, Jingwei Li, Haiyan Huang.

**Formal Analysis:** Tianjie Li, Jingwei Li, Leyang Wang, Haiyan Huang.

**Funding Acquisition:** Qiang Wu.

**Investigation:** Tianjie Li.

**Methodology:** Tianjie Li.

**Resources:** Qiang Wu.

**Writing – Original Draft Preparation:** Tianjie Li.

**Writing – Review & Editing:** Leyang Wang, Jingwei Li, Haiyan Huang, Qiang Wu.

