## Supplemental FigureS1-6 for "Wiz regulates clustered protocadherin genes by restricting CTCF/cohesin loop extrusion in a genomic-distance biased manner"

bioRχiv

**The PDF file includes:**

**Supplementary Figures S1 to S6**

**S1 Fig.** Model architectures and predicted zinc finger distribution.

**S2 Fig.** COP prediction of C2H2-ZFP occupancy at the *cPcdh* pCBS elements.

**S3 Fig.** Generation of N2a single-cell clones with *Wiz* Myc-tagged or deleted.

**S4 Fig.** Increased enrichment of active chromatin marks of H3K4me3 and H3K27ac at *cPcdh* regulatory elements upon *Wiz* deletion.

**S5 Fig.** Generation of *Wiz* conditional knockout mice via pronuclei injection.

**S6 Fig.** Repressive chromatin marks of H3K9 mono-, bi-, and tri-methylations show no obvious alteration upon *Wiz* deletion.

**Captions for Tables S1 to S4**

**S1 Table.** Oligonucleotides used in this study.

**S2 Table.** Datasets used in this study.

**S3 Table.** COP prediction of C2H2-ZFPs' occupancy at the pCBS elements of the *Pcdh* clusters.

**S4 Table.** Performance comparison of COP and common machine learning classifiers.

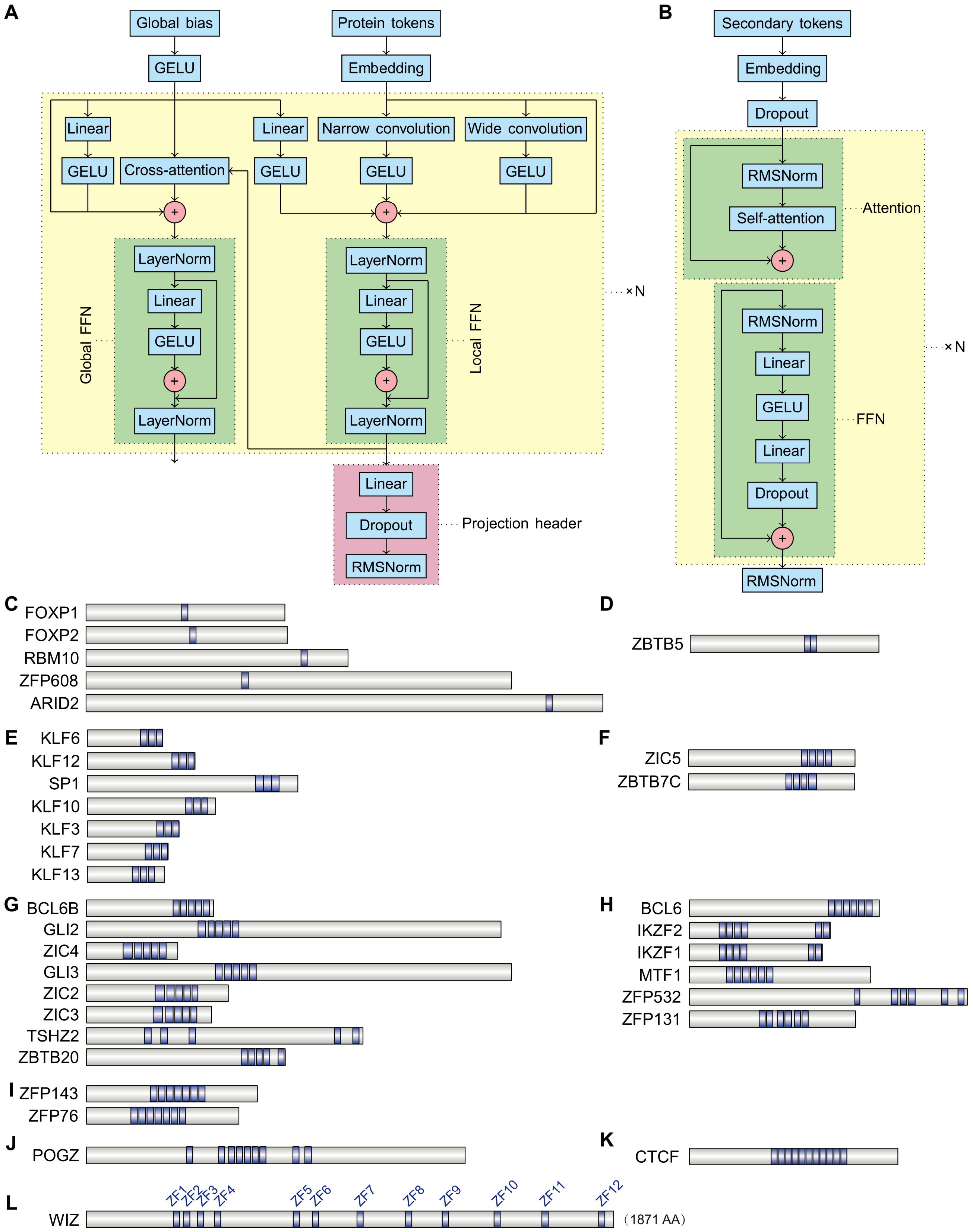


**S1 Fig. Model architectures and predicted zinc finger distribution.** (**A**) Architecture of the ProteinBERT model for deep learning on protein sequences. Protein residue-wise features are processed by narrow and wide convolutions. Cross-attention operates between a single global embedding and multiple residue embeddings, resulting in linear complexity with protein length. Bidirectional information flow between local and global representations allows residues to depend on each other. A projection head ensures output dimensionality consistent with COP. GELU, Gaussian error linear unit. (**B**) Architecture of the protein secondary structure encoder, implemented as a standard transformer. RMS normalization was used in place of layer normalization and applied prior to the attention module. (**C-L**) COP predicts 34 C2H2-ZFP members potentially occupy all 54 *cPcdh* promoter CBS elements, with 5 contain single C2H2 zinc finger (ZF) domain (**C**), 1 contains two C2H2-ZFs (**D**), 7 contain three C2H2-ZFs (**E**), 2 contain four C2H2-ZFs (**F**), 8 contain five C2H2-ZFs (**G**), 6 contain six C2H2-ZFs (**H**), 2 contain seven C2H2-ZFs (**I**), 1 contains nine C2H2-ZFs (**J**), 1 contains ten C2H2-ZFs (**K**), and 1 contains twelve C2H2-ZFs (**L**). Blue box indicates C2H2-ZF domain.

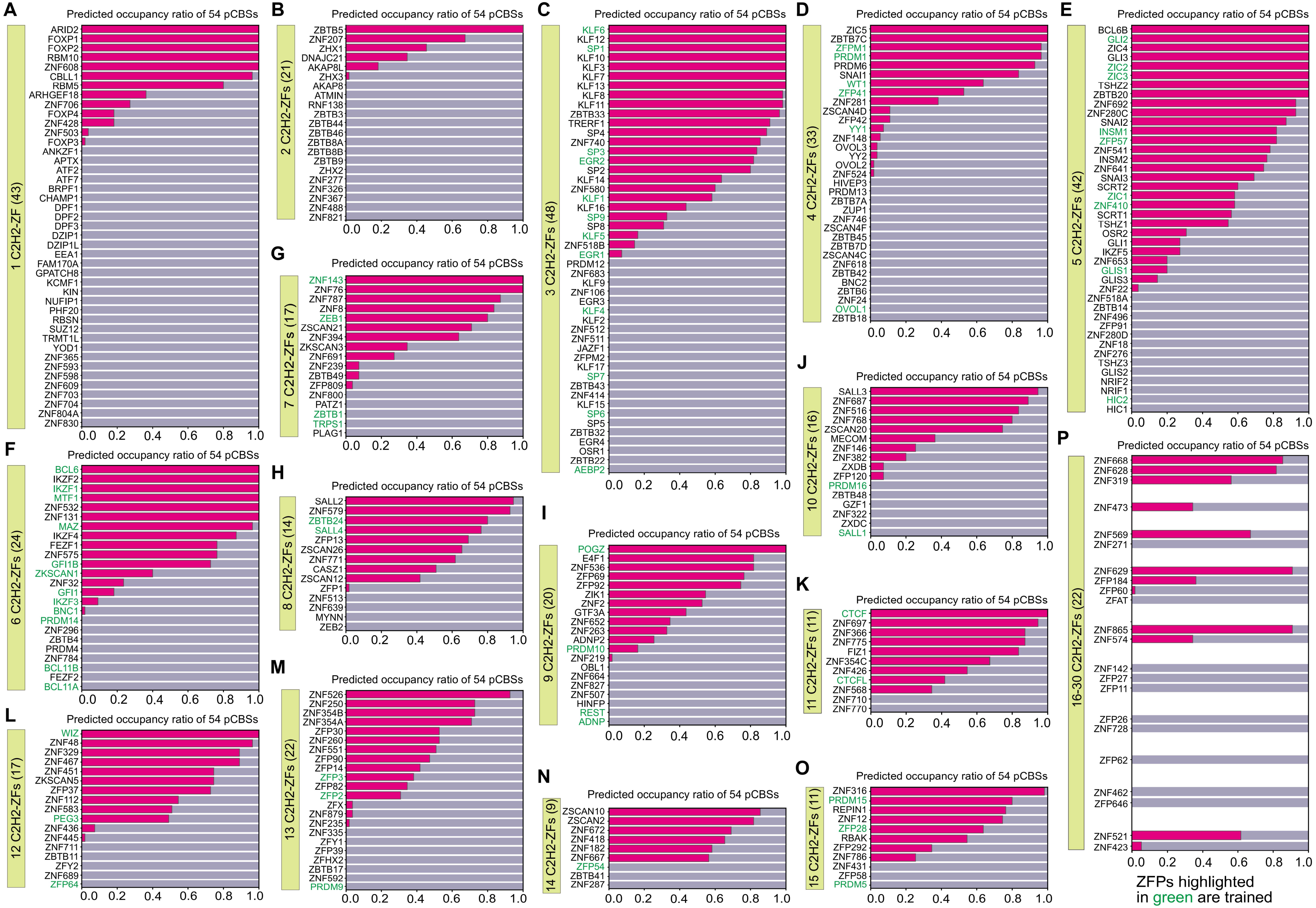

**S2 Fig. COP prediction of C2H2-ZFP occupancy at the *cPcdh* pCBS elements.** (**A-P**) COP-predicted occupancy ratio at the 54 *cPcdh* pCBS elements for each C2H2-ZFP containing single C2H2-ZF domain (**A**), as well as containing 2 (**B**), 3 (**C**), 4 (**D**), 5 (**E**), 6 (**F**), 7 (**G**), 8 (**H**), 9 (**I**), 10 (**J**), 11 (**K**), 12 (**L**), 13 (**M**), 14 (**N**), 15 (**O**), or 16-30 (**P**) C2H2-ZFs. C2H2-ZFPs are colored in green for trained or black for untrained. The protein numbers for each ZFP group with 1-30 ZFs are indicated in parentheses.

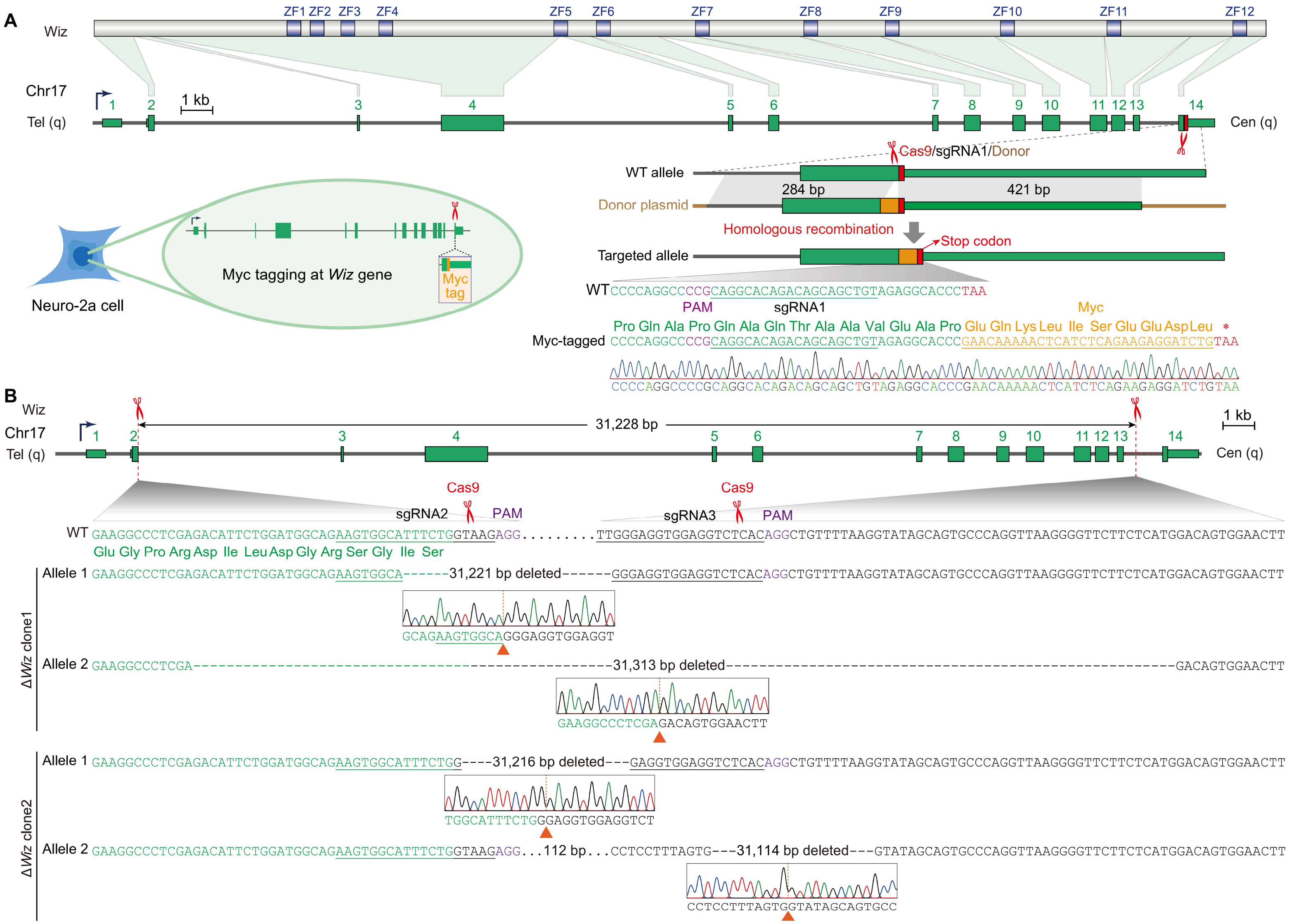


**S3 Fig. Generation of N2a single-cell clones with *Wiz* Myc-tagged or deleted.** (**A**) Generation of N2a single-cell clones with endogenous *Wiz* Myc-tagged at C-terminus. A Myc-coding sequence was inserted immediately upstream of the stop codon (TAA) of the endogenous *Wiz* gene via single-sgRNA-guided Cas9 cleavage followed by DNA template-mediated homology-directed repair (HDR). C-terminal Myc tagging was confirmed in single-cell clone by genotyping with Sanger sequencing. The inserted Myc tag is indicated in orange. (**B**) Generation of *Wiz*-knockout N2a single-cell clones via CRISPR/Cas9-mediated DNA fragment deletion. Two sgRNAs were designed to program Cas9 cleavage within introns 2 and 13 of the *Wiz* gene, respectively, enabling excision of the intervening genomic fragment. Genotyping of *Wiz*-knockout (Δ*Wiz*) single-cell clones by Sanger sequencing confirmed large, targeted deletions. Junction sequence analyses revealed substantial allelic heterogeneity, attributable to variable end resection and error-prone non-homologous end joining (NHEJ) repair at the double-strand breaks (DSBs) on individual alleles.

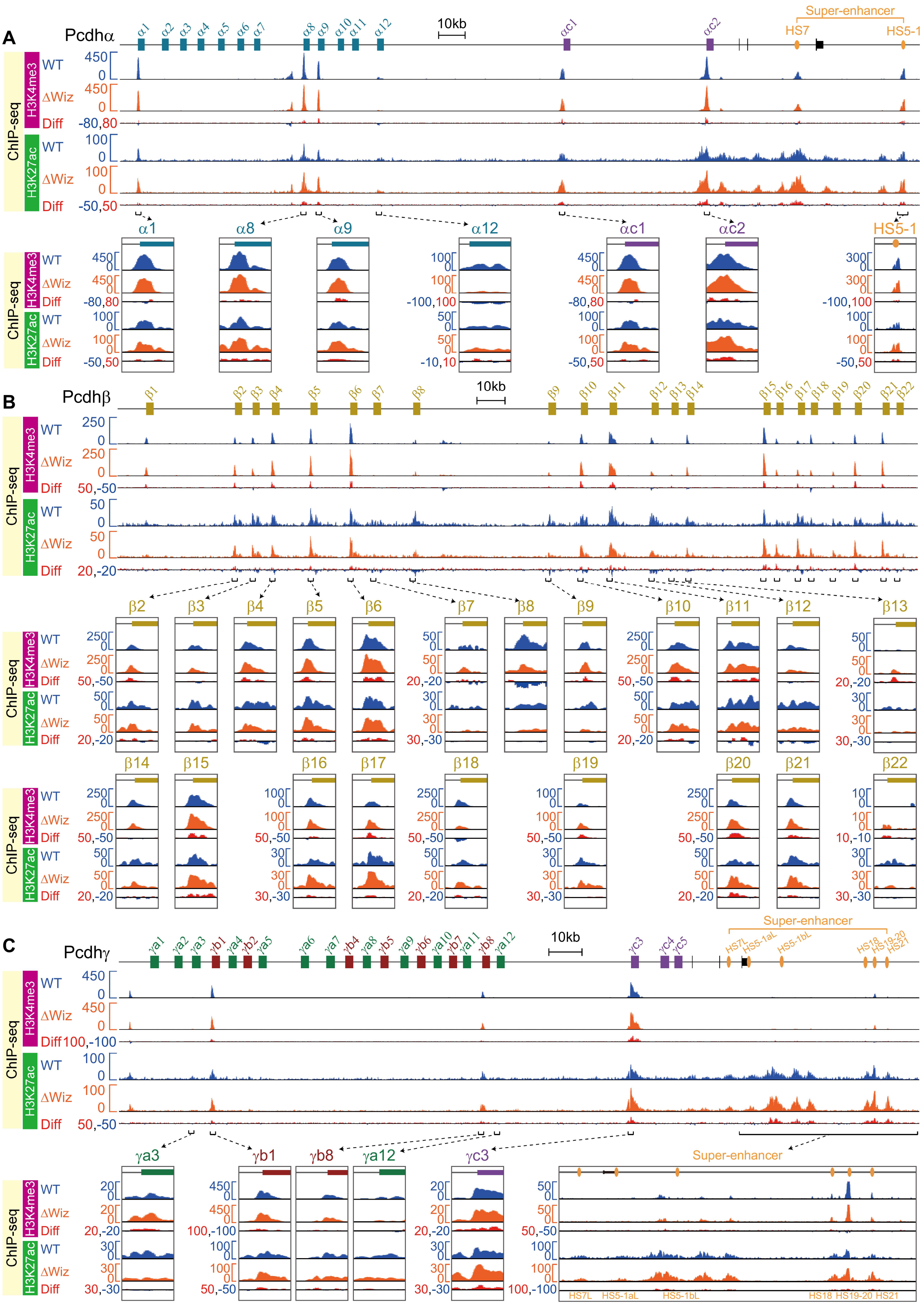


**S4 Fig. Increased enrichment of active chromatin marks of H3K4me3 and H3K27ac at *cPcdh* regulatory elements upon *Wiz* deletion.** (**A-C**) ChIP-seq profiles of H3K4me3 and H3K27ac at the *Pcdh* $\alpha$ (**A**), $\beta$ (**B**), and $\gamma$ (**C**) gene clusters in Δ*Wiz* N2a single-cell clones compared to wild-type (WT) control clone. Close-up views of promoter and enhancer regions reveal increased enrichments of both H3K4me3 and H3K27ac, marks of active chromatin, upon *Wiz* deletion.

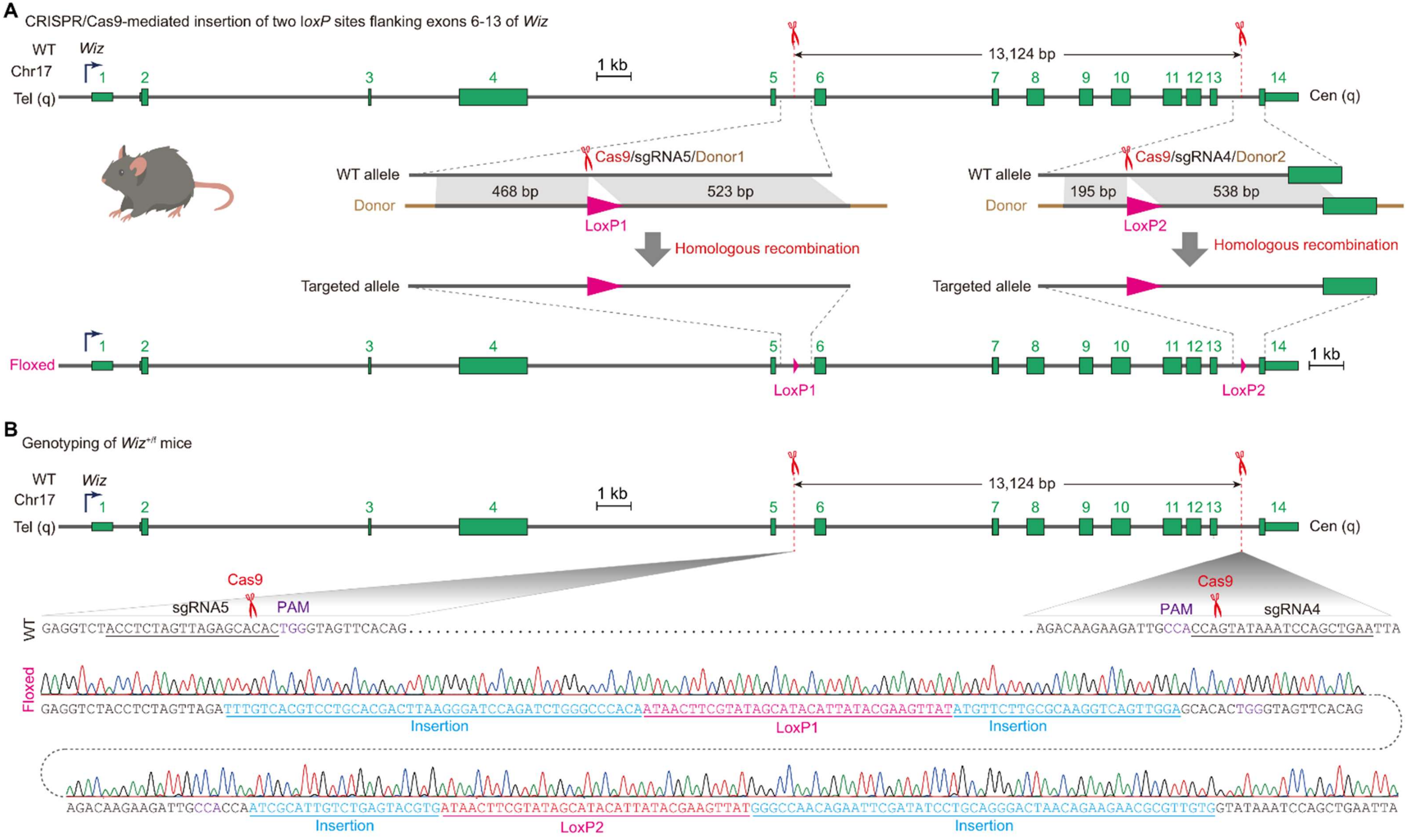


**S5 Fig. Generation of *Wiz* conditional knockout mice via pronuclei injection.** (**A**) Schematic of CRISPR/Cas9-mediated homologous recombination (HR) for generating conditional *Wiz* knockout mouse model. Two *loxP* sites were precisely inserted into introns 5 and 13 of the *Wiz* gene via Cas9-induced double-strand breaks (DSBs), guided by single sgRNAs and repaired using donor DNA templates with the loxP sites. (**B**) Genotyping of the homologous *Wiz*-floxed ($Wiz^{f/f}$) mouse strain by Sanger sequencing confirming targeted insertion of *loxP* sites flanking exons 6-13 of the *Wiz* gene.

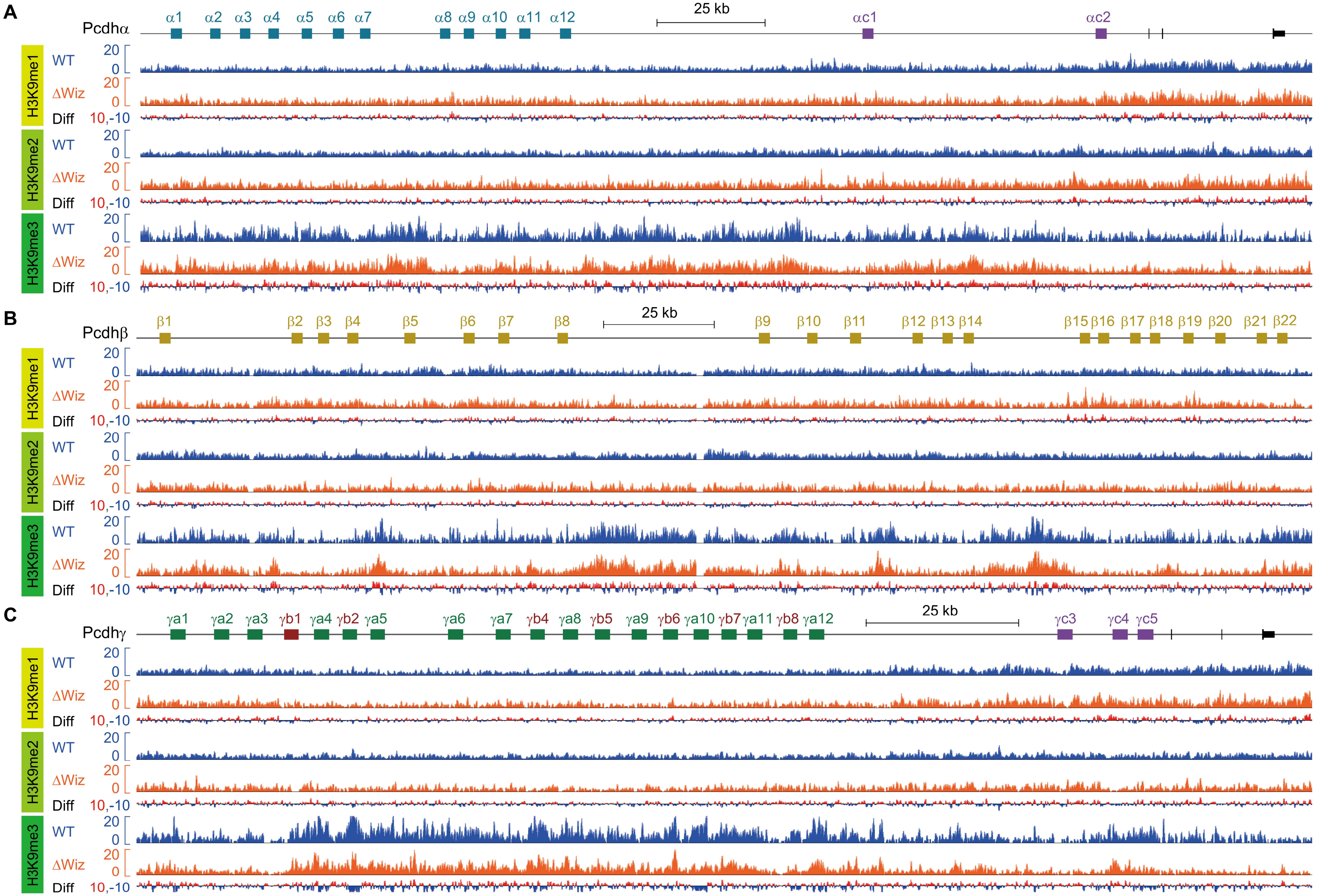


**S6 Fig. Repressive chromatin marks of H3K9 mono-, bi-, and tri-methylations show no obvious alteration upon *Wiz* deletion.** (**A-C**) ChIP-seq profiles showing no obvious alterations of repressive chromatin marks of H3K9me1, H3K9me2, and H3K9me3 across *Pcdhα* (**A**), *β* (**B**), and *γ* (**C**) clusters.
